# ATR, a DNA damage kinase, modulates DNA replication timing in *Leishmania major*

**DOI:** 10.1101/2025.01.13.632695

**Authors:** Gabriel L. A. da Silva, Jeziel D. Damasceno, Jennifer A. Black, Richard McCulloch, Luiz R. O. Tosi

## Abstract

All cells possess mechanisms to maintain and replicate their genomes, whose integrity and transmission are constantly challenged by DNA damage and replication impediments. In eukaryotes, the protein kinase Ataxia-Telangiectasia and Rad3-related (ATR), a member of the phosphatidylinositol 3-kinase-like family acts as a master regulator of the eukaryotic response to DNA injuries, ensuring DNA replication completion and genome stability. Here we aimed to investigate the functional relevance of the ATR homolog in the DNA metabolism of *Leishmania major*, a protozoan parasite with a remarkably plastic genome. CRISPR/cas9 genome editing was used to generate a Myc-tagged ATR cell line (mycATR), and a Myc-tagged C-terminal knockout of ATR (mycATRΔC-/-). We show that the nuclear localisation of ATR depends upon its C-terminus. Moreover, its deletion results in single-stranded DNA accumulation, impaired cell cycle control, increased levels of DNA damage, and delayed DNA replication restart after replication stress. In addition, we show that ATR plays a key role in maintaining *L. major’s* unusual DNA replication program, where larger chromosomes duplicate later than smaller chromosomes. Our data reveals loss of the ATR C-terminus promotes the accumulation of replication signal around replicative stress fragile sites, which are enriched in larger chromosomes. Finally, we show that these alterations to the DNA replication program promote chromosome instability. In summary, our work shows that ATR acts to moderate DNA replication timing thus limiting the plasticity of the *Leishmania* genome.

## Introduction

DNA stability is constantly challenged by a range of stressors that can cause a diversity of DNA lesions(1,2). To tackle all possible lesion types and guarantee genome integrity, cells have evolved a range of pathways, collectively known as the DNA Damage Response (DDR), which when activated, lead to damage recognition, signalling and, ultimately repair (2–4). At the pinnacle of the eukaryotic DDR are three protein kinases that act to recognise and signal DNA damage, trigger the activation of the appropriate repair response: ATM (Ataxia-Telangiectasia Mutated) and DNA-PKcs (DNA protein kinase catalytic subunit) are recruited primarily to DNA double strand breaks (DSBs), whereas ATR (Ataxia-Telangiectasia and Rad3-related) primarily responds to the accumulation of single-stranded DNA (ssDNA) during DNA replication (3–5). Stalling or slowing of the replisome leads to ssDNA accumulates during DNA replication (6). Exposed ssDNA is protected by the RPA complex, limiting degradation by nucleases (6,7). The resulting ssDNA-RPA complex, along with 5’-ended ssDNA-dsDNA junctions formed at the stalled replication fork, serves as a platform for the recruitment of ATR-interacting protein (ATRIP), TOPBP1 (Topoisomerase II-binding protein 1) and the heterotrimeric 9-1-1 checkpoint complex (Rad9-Rad1-Hus1) (8–10). ATRIP recruits ATR to the damage site, while TOPBP1 and 9-1-1 complex are responsible for the stabilisation and full activation of ATR (10–13). In addition to these initial factors, in some organisms ETAA1 also activates ATR by directly interacting with RPA (14,15). Once activated, ATR coordinates a complex response that safeguards the replication fork to preserve DNA integrity (14). Instrumental to this response is the effector kinase CHK1, which is activated upon phosphorylation by ATR (16,17). ATR-driven CHK1 activation functions to arrest the cell cycle, suppress origin firing, stabilize replication forks, and promote fork repair and DNA replication restart (18,19). By coordinating cell cycle arrest with fork stabilization and recovery, ATR and CHK1 prevent cells from entering mitosis when DNA replication is compromised (19,20). Most of our understanding of ATR’s-activities come from studying mammalian and yeast cells (21–24). Far less research has examined how the ATR pathway operates in eukaryotic pathogens(25). *Trypanosoma brucei*, *Trypanosoma cruzi* and *Leishmania spp.* are related protozoan parasites, part of a wider grouping known as trypanosomatids. These protozoa cause vector-bone diseases affecting millions of people every year in tropical and sub-tropical regions of the globe, though recent cases of Chagas disease and leishmaniasis are emerging across Europe and North America (26–29). All three pathogens are notable for their complex adaptive changes to their genome composition to survive different environments throughout their life cycles (30–32). For instance, *T. brucei* and *T. cruzi* display elevated levels of recombination amongst large, variable gene families that encode cell surface proteins required for evasion of the host immune system(33–35). *Leishmania* lack such variable gene families; however, genome plasticity is remarkably widespread, with both natural isolates and laboratory strains showing chromosomal aneuploidy and gene copy number variation (CNV). Such alterations to their genome composition are commonly associated with drug resistance emergence and environmental adaptations (31,32,36–38).

Several studies have linked DDR catalytic factors to mutation and Copy Number Variation (CNV) (39–41), but fewer have examined how damage signalling may influence genome stability (25). To date, there has been no investigation into whether or not *Leishmania* DDR kinases modulate genome stability. Studies in *T. brucei* show that ATR loss has a distinct impact depending upon the life cycle stage: in insect procyclic stages, RNAi-mediated depletion of ATR leads to a modest reduction in cell proliferation and cell cycle progression following ionizing radiation exposure (42), while in mammalian-infective cells ATR depletion is lethal and disrupts VSG expression (42,43). In *Leishmania*, initial studies investigating the potential functions of ATR used human-specific ATR inhibitors, reporting a reduction in cell proliferation, increased sensitivity to oxidative stress, and an accumulation of ssDNA (44,45). Gene knockout experiments in *L. mexicana* suggested that ATR is not essential for cell survival *in vitro* (46), and our recent work suggests that ATR deficiency in *L. major* exacerbates replication stress (47). In this study, we used CRISPR/Cas9 genome editing to generate an ATR mutant cell line. Although a complete KO of the *ATR* gene could not be achieved in *L. major*, we show that loss of the predicted kinase domain, located within the C-terminus of the kinase, disrupts nuclear localization and is tolerated. Using this mutant, we demonstrate that ATR is involved in regulating the unusual chromosome size-dependent timing of *Leishmania* DNA replication, and that disrupting this process leads to compromised genome stability and variation.

## Results

### Generation of ATR mutants lacking the C-terminal kinase domain in *Leishmania major*

To investigate the roles of ATR in *L. major*, we sought to genetically modify the *ATR* locus (LmjF.32.1460) for two purposes. To first localise the protein, we used CRISPR-Cas9 to generate cells expressing full length ATR tagged at its N-terminus by inserting mNeonGreen (mNG) and three copies of Myc tag(48,49) upstream of the *ATR* ORF (Figure 1A). These cells are referred to as mycATR. Second, to assess ATR function, we aimed to delete the ATR ORF. However, all attempts to remove the entire *ATR* ORF using CRISPR/Cas9 were unsuccessful, suggesting ATR is essential in the promastigote forms of *L. major*. Therefore, in our mycATR cells we opted to delete a 3,444bp region from the C-terminus of *ATR* (from 6180 to 9624 bp), containing the predicted kinase domain, generating the cell line mycATRΔC-/- (Figure 1A).

**Figure 1.**
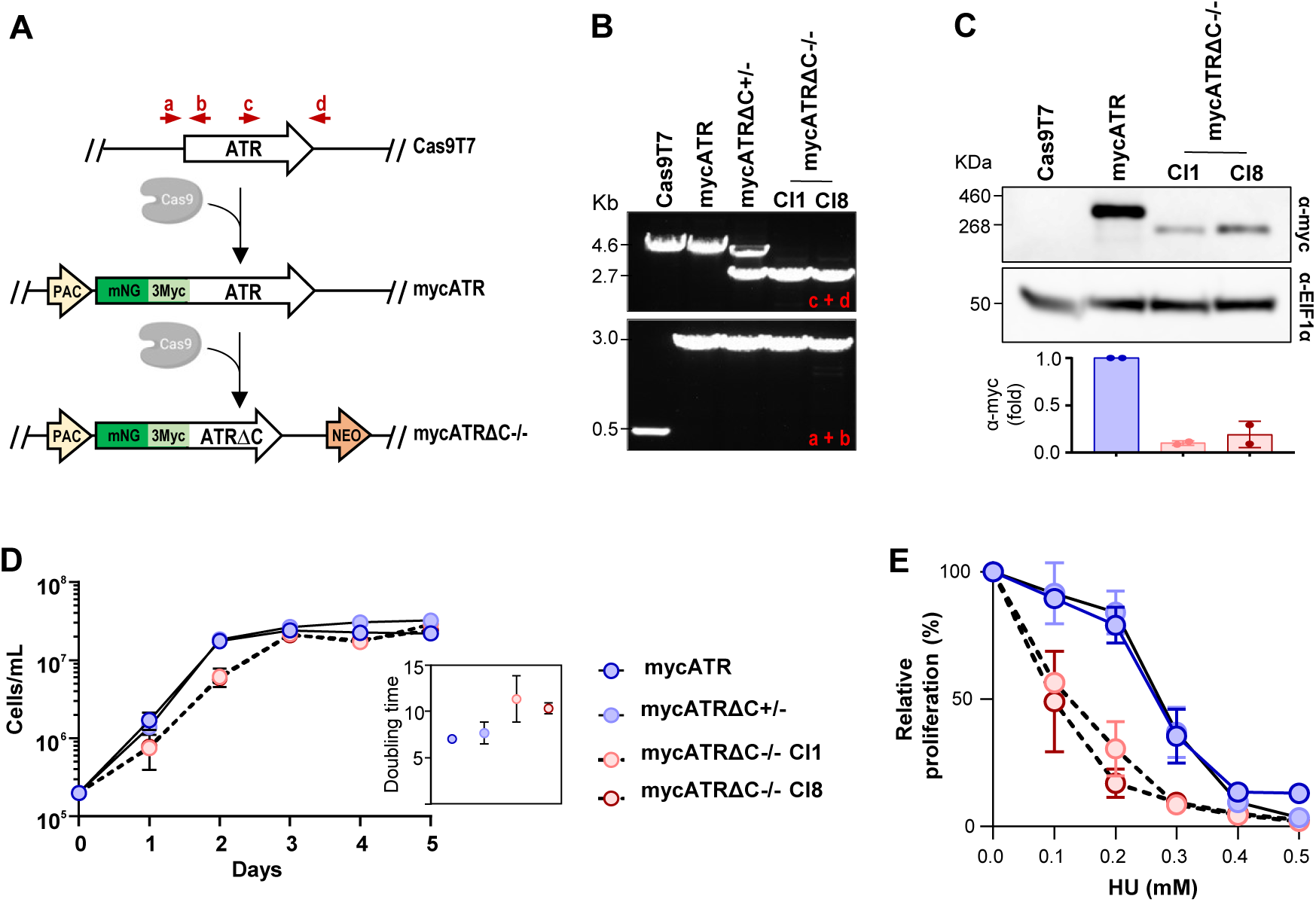
Deletion of the C-terminus of *L. major* ATR alters the protein’s abundance and sensitivity to replication stress. (**A**) Schematic representation showing a cell line expressing Cas9, which was used to add a tag (3xmyc + NeonGreen) N-terminally tio the ATR kinase (LmjF.32.1460), generating mycATR cell line. The predicted kinase domain present in the C-terminal region (∼6220 – 9624 pb) of the mycATR cell line was replaced by a selectable marker (NEO), generating the mycATRΔC-/- cell line; red arrows represents primers used to detect these alterations. (**B**) PCR analysis from mycATR, mycATRΔC+/- and mycATRΔC-/- cell lines, indicating the addition of the tags in two alleles of both cell lines (a + b) and the deletion of the C-terminal region (c + d) in both alleles of mycATRΔC-/- cells. (**C**) Western analysis of whole cell extracts of indicated cell lines in exponential growth; extracts were probed with α-myc antiserum, and α-EIF1α was used as a loading control (predicted protein sizes are indicated, kDa). Quantification of the band from the western blot showed in (C) using ImageJ software; statistical analysis was performed using Prism (Error bars ± SD; n = 2 experiments, * p<0.05; ** p<0.005; *** p<0.001; **** p<0.0001. Unpaired t-test). (**D**) Growth curve of the indicated cell lines cultivated in HOMEM medium; cells were seeded at ∼2x105 cells.ml-1 at day 0; growth profile was evaluated after cells were kept in culture for 5 days and the cell density was assessed every 24 hours (Error bars ± SD; n = 2 experiments). (**E**) Cell survivl curve of indicated cell lines cultivate in HOMEM medium; cells were seeded at ∼2x105 cells.ml-1 and left untreated or treated with the indicated concentrations of Hydroxyurea HU (mM); after 96 hours the cell density was assessed and the percentage was calculated relative to the density of untreated cells.

PCR amplification of the N- or C-termini of the *ATR* locus revealed that a single round of antibiotic selection was successful to tag the N-terminus of ATR and to delete the C-terminal coding domain of both alleles (Figure 1B). We independently generated two such deletion clones, cl1 and cl8. Additionally, we recovered a clone in which only one allele of *ATR* was truncated, referred to as mycATRΔC+/-. Expression of tagged ATR in the mycATR and mycATRΔC-/- cells was assessed by western blotting, revealing bands of expected: full-length tagged ATR migrated just below a 460 kDa marker (predicted size, 383.31 kDa), while the C-terminally truncated version appeared at approximately 260 kDa (predicted size, 256.42 KDa; Figure 1C). Quantification of myc signal on the blot further showed that the C-terminal deletion reduced total ATR protein levels, indicating that this region could be important for ATR stabilization and/or expression (Figure 1C).

Next, we examined the growth profile of mycATR, mycATRΔC-/-, and mycATRΔC+/- cell lines (Figure 1D), comparing their growth to that of parental *L. major* CC1 and Cas9-T7 cells (Figure S1A). The results showed that loss of one *ATR* allele or N-terminal tagging of full-length ATR did not affect population doubling (Figure 1D and S1A). In contrast, proliferation of the mycATRΔC-/- cells was reduced compared to mycATR and mycATRΔC+/- cells, resulting in an increase in doubling time (mycATRΔC-/- clones, ∼11 and 10 hours, versus ∼7 and 8 hours for mycATR and mycATRΔC+/- cells, respectively; Figure 1D). Despite this difference in proliferation, FACS analysis of DNA synthesis, assessed by IdU incorporation, showed similar percentages of replicating cells in the mycATR and mycATRΔC+/- populations (S1B). Also, all mutant clones reached the same maximum density as the mycATR and mycATRΔC+/- cells (Figure 1D). We next assessed the sensitivity of mycATR, mycATRΔC+/- and mycATRΔC-/- cells to hydroxyurea (HU), a reversible inhibitor of ribonuceotide reductase. Cells were treated with increasing concentrations of HU or left untreated. After 96 hours of growth, we measured cell density in the HU-treated populations relative to untreated controls. The results showed that mycATRΔC-/- cells grew slower or died more quickly in the presence of HU compared with both mycATR and mycATRΔC+/- cells (Figure 1E). Overall, these findings indicate that ATR C-terminal deletion alters promastigote growth dynamics and increases sensitivity to HU-induced replication stress, highlighting a role for ATR during the replication stress of *L. major*.

### The C-terminus of ATR is vital for accumulation of the protein in the nucleus

In other eukaryotes, including *T. brucei*, ATR has been shown to localize to the nucleus (43,50,51). However, the subcellular localization of ATR in *Leishmania* species remains unknown from available data on Leishtag.org). To investigate the subcellular location of ATR in *L. major* promastigotes and ask if deletion of the C-terminal plays a role in ATR localisation, we performed indirect immunofluorescence (IFA) on formaldehyde-fixed mycATR and mycATRΔC-/- cells using anti-myc antiserum (α-myc). Images were captured using structured illumination super-resolution microscopy (SR-SIM), allowing for high-resolution visualization of ATR localization (Figure 2A). IFA revealed accumulation of α-myc signal within the nuclear compartment of mycATR cells. In contrast, mycATRΔC-/- cells showed little nuclear localization, with α-myc signal distributed throughout the cell body, including signal throughout the cytoplasm (Figure 2A). Quantification of the nuclear myc signal confirmed a significant decrease in nuclear myc signal intensity in mycATRΔC-/- cells compared with mycATR cells (Figure 2B), suggesting that deletion of the C-terminus disrupts nuclear localisation of ATR. In support of a nuclear localisation for ATR directed by the C-terminus, we identified a putative nuclear location signal (NLS) within the deleted region of the ATR ORF (amino acids 2922 to 2932) (Figure S2A).

**Figure 2.**
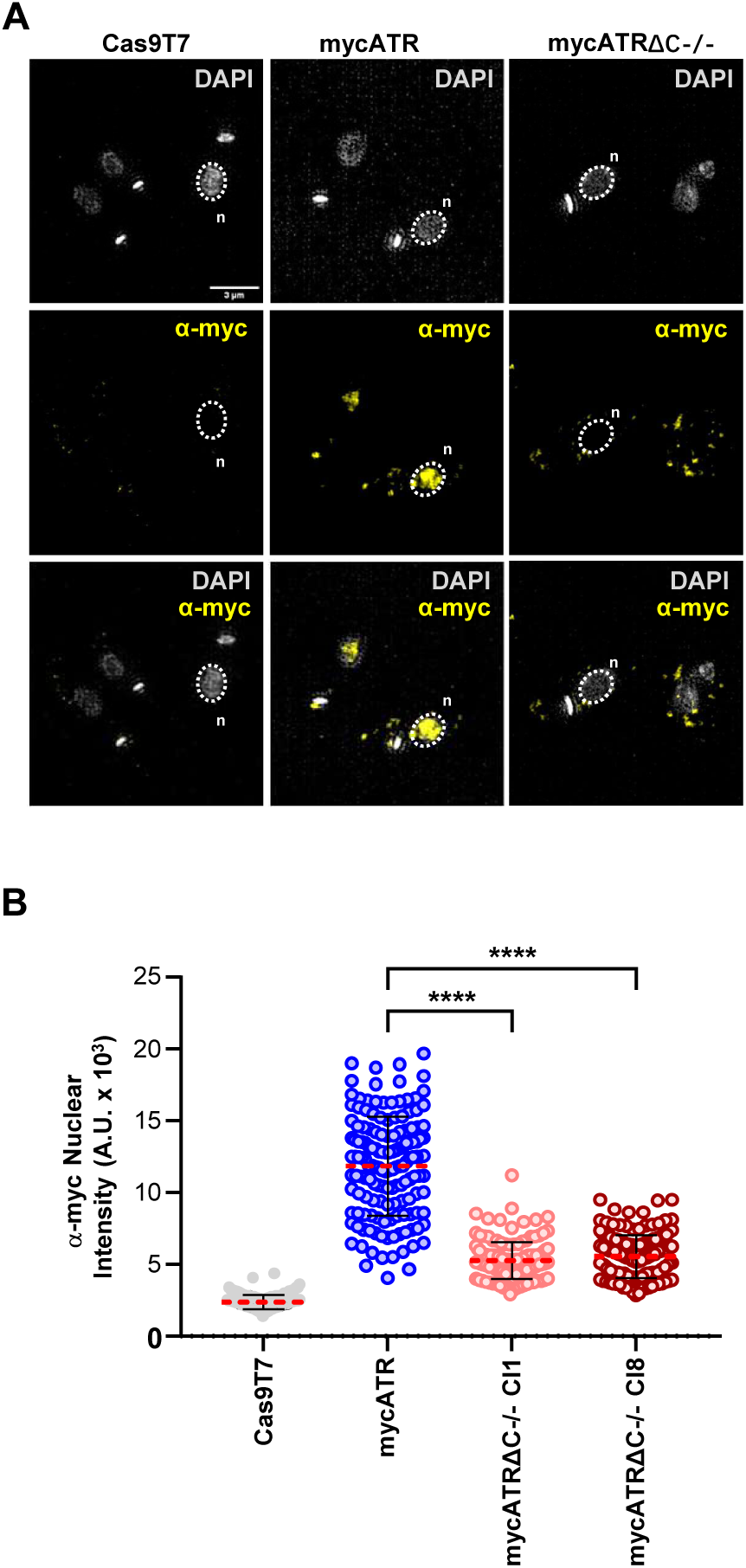
Subcellular location of *L. major* ATR kinase depends on its C-terminus. (**A**) Representative images of the sub-cellular localization of myc signal, which is added N-terminally to ATR in mycATR and mycATRΔC-/- cells; scale bar = 3 μm. Images were captured on an Elyra super resolution microscope (Zeiss). Nuclear (n) and Kinetoplast (k) DNA are shown in gray; and the myc signal in yellow. (**B**) Quantification of the myc signal around the nucleus. An area was drawn around ≥ 100 nuclei based on the DAPI stain and the myc signal was measured using ImageJ software, Cas9T7 cells were use as negative control (Error bars ± SD; n = 2 experiments, **** P<0.0001. Unpaired t-test).

To further validate the subcellular location of ATR, we performed live cell microscopy using mycATR cells, but this time detecting mNG signal. (Figure). Additionally, we tagged ATR with just three copies of Myc (3xMycATR) to ask if mNG tagging altered protein localisation. Both approaches confirmed the accumulation of ATR within the nuclear compartment (Figure S2B and S2C). Altogether, we suggest that the growth and HU sensitivity phenotypes associated with the loss of the C-terminus of ATR may arise from a combination of lost C-terminal functions, reduced protein abundance and impaired accumulation in the nucleus.

### Replication stress leads to ssDNA accumulation and cell cycle defects, which impairs replication restart in mycATRΔC-/- cells

The primary substrate of ATR activation is elongated tracks of ssDNA (13,52). In ATR’s absence, ssDNA accumulates in human cells under replication stress conditions (50,53). Given that even partial deficiency of ATR increases ssDNA levels in *L. major* (47), we investigated whether deletion of the C-terminus of ATR leads to ssDNA accumulation. To assess this, we labelled DNA by growing mycATR and mycATRΔC-/- cells in medium containing IdU, a thymidine analogue. Cells were then washed and cultivated in the presence or absence of 5 mM of HU for 5 hours. Cells were fixed and stained with α-BrdU antiserum to detect the incorporated IdU, with α-BrdU signal reflecting accumulation of ssDNA, since DNA was not denatured.

Microscopy images showed a clear increase of nuclear IdU signal in HU-treated mycATRΔC-/- cells compared with untreated control, and with both treated and untreated mycATR cells (Figure 3A). Quantification of the signal intensity confirmed a significant increase in the HU-treated mycATRΔC-/- cells (Figure 3B), indicating that loss of ATR leads to the accumulation of ssDNA in *L. major*.

**Figure 3.**
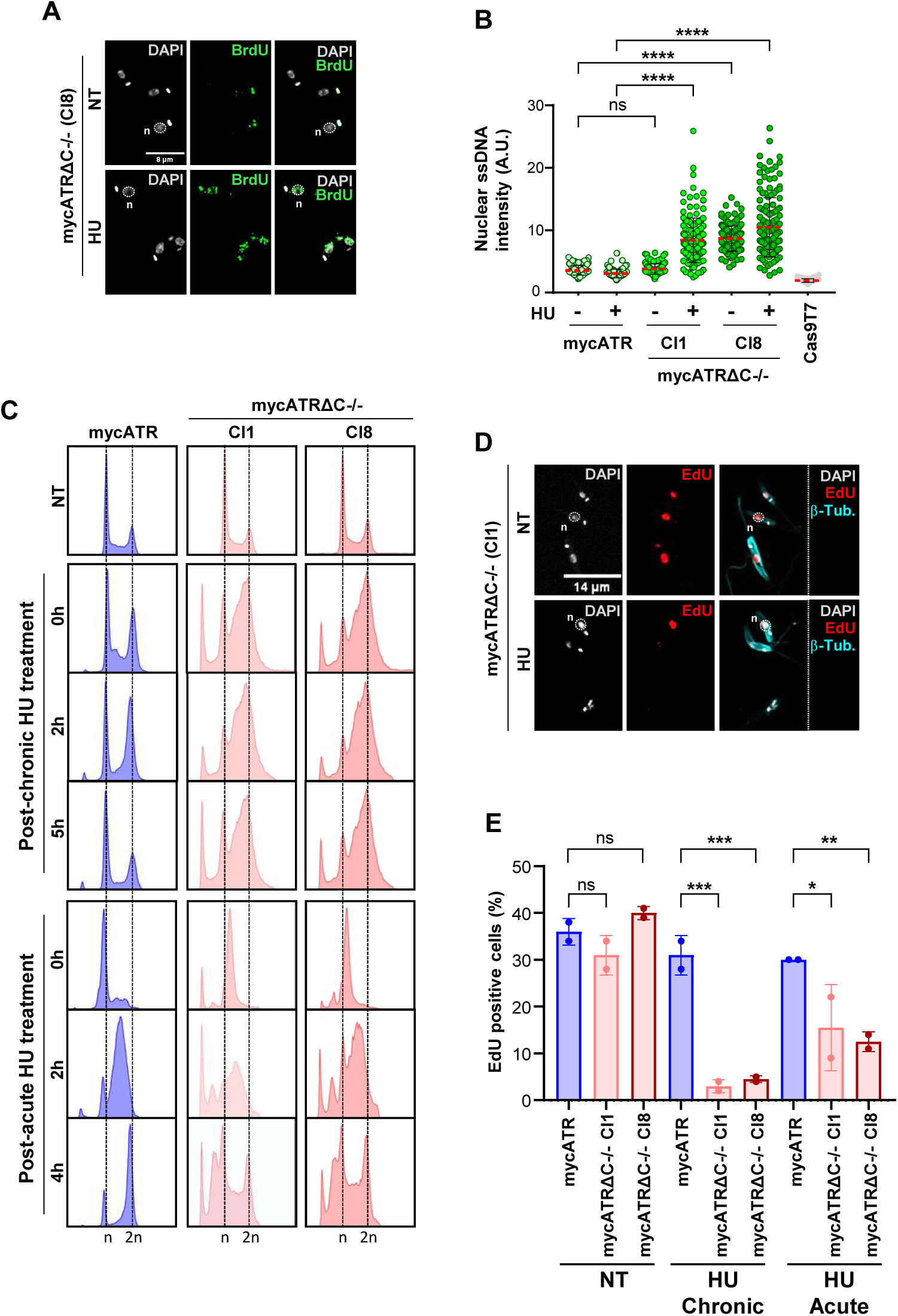
*L. major* ATR protects the cell cycle after replication stress by preventing ssDNA accumulation and promoting DNA replication restart. (**A**) Representative images acquired in LSM880 Zeiss confocal microscope from Z stack images of mycATRΔC-/- (cl8) untreated (NT) or treated with 5mM of HU for 5 hours. Cells were fixed with 3% formaldehyde and the stained with α-BrdU (ssDNA, green), and the genomic DNA stained with DAPI (gray). Bar scale = 8 um. (**B**) Quantification of the ssDNA signal in the nucleus of mycATR and mycATRΔC-/- cells treated or not with 5 mM of HU for 5 hours; Cas9T7 cells without α-BrdU were use as negative control. An area was drawn around 150 nuclei based on the DAPI stain and the ssDNA signal was measured using ImageJ software (Error bars ± SD; n = 2 experiments, ** p<0.005; *** p<0.001; **** p<0.0001. 2way ANOVA). (**C**) mycATR and mycATRΔC-/- cells were left untreated (NT) or treated with 0.5 mM of Hydroxyurea (HU) for 20 hours (Chronic) or treated with 5 mM of Hydroxyurea (HU) for 6 hours (Acute), washed and resuspended in fresh media, and samples collected i0, 2, 5 hours after chronic treatment and 0, 2, 4 hours after acute treatment. Cells were fixed with methanol and the DNA stained with propidium iodate, followed by flow cytometer analysis. (**D**) Representative images acquired in LSM880 Zeiss confocal microscope from Z stack images of mycATRΔC-/- (cl1) where cells were pulsed with 150 uM of EdU for 30 minutes in untreated condition or after release from chronic (5 hours) or acute (4 hours) treatment. Cells were fixed with 3% formaldehyde and a Click-IT reaction was used to stain cells that had uptaken EdU during 30 min (red); genomic DNA stained with DAPI (gray) and α-tubulin was used to stain the cytoskeleton (cyan). Bar scale = 14 um. (**E**) Percentage of EdU positive mycATR or mycATRΔC-/- cells in each condition was calculated by the number of stained cells by the number of cells present in each sample (100 – 300 cells; error bars ± SD; n = 2 experiments, * p<0.05; ** p<0.005; *** p<0.001, One-way ANOVA).

Considering that one of the primary functions of ATR, once activated by ssDNA accumulation, is cell cycle regulation (54–56), we next examined the cell cycle profiles of mycATR and mycATRΔC-/- cells by flow cytometry with and without HU exposure in exponentially growing cells. We selected two different conditions to examine the effect of ATR C-terminal loss: chronic (0.5 mM of HU for 20 h) or acute replication stress challenge (5 mM of HU for 5 h). Cells were collected and fixed after being cultured in fresh media for 0, 2 and 5 h following chronic HU treatment, or for 0, 2 and 4 h after acute HU treatment. The FACS profile of propidium iodide-stained DNA of mycATR cells showed that chronic treatment did not completely block cell cycle progression; rather the progression through S phase was slowed. Specifically, there was an initial accumulation of cells in early S phase at 0 h, followed by an accumulation in mid/late S phase at 2 h after HU release. By 5 hours, the cell cycle profile had returned to levels like those seen in untreated cells (Figure 3C). The higher concentration of HU during the acute treatment induced a cell cycle stall in the mycATR cells at the G1/S boundary (0 h, Figure 3C). After release from HU, mycATR cells progressed through S phase into G2/M at 2 and 4 h, respectively (Figure 3C). In contrast, both HU treatments had a distinct effect on cell cycle progression in mycATRΔC-/- cells. After chronic HU exposure, we detected an increased accumulation of mycATRΔC-/- cells in S phase at 0 h (Figure 3C), which did not obviously alter at either 2 or 5 h following HU release.

Moreover, we observed an increase in the number of sub-G1 cells (<G1) following chronic HU exposure in the absence of the C-terminus of ATR (Figure 3C). After acute HU treatment, mycATRΔC-/- cells synchronised at the G1/S boundary (0 h, Figure 3C) and progressed through S phase to G2/M between 2 and 4 h, but with a substantial rise in the <G1 population (Figure 3C). Taken together, these data suggest that deleting the C-terminus of ATR leads to cell cycle defects after HU exposure. Chronic treatment with 0.5 mM HU slows cell cycle progression of mutant cells and induces cells with aberrant DNA content, while acute treatment with 5 mM HU allows cell cycle progression after G1/S synchronisation, but it is also associated with the generation of pronounced levels of aberrant cells. Overall, mycATRΔC-/- cells leads to aberrant cell cycle progression under increased levels of DNA replication stress.

The ability of stalled DNA polymerases to restart DNA synthesis after removal or bypass of a block in DNA replication elongation is an indication of replication fork stabilization(57–59). To further examine the effects of replication stress in the absence of ATR’s C-terminus, we investigated the percentage of mycATR and mycATRΔC-/- cells capable of restarting DNA synthesis after replication stress. Untreated cells, cells 5 h after chronic HU exposure, or 4 h after release from G1/S arrest by acute HU exposure were pulsed with 150 uM EdU for 30 min. Cells were then fixed and a Click-It reaction was performed to detect cells that incorporate the nucleotide analogue (as a proxy for DNA synthesis; Figure 3D). The percentage of EdU positive cells were identified by microscopy and calculated as a proportion of the total of cells population (Figure 3E). In untreated conditions, we detected no difference in the percentage of EdU-positive mycATR and mycATRΔC-/- cells (∼30% for both; Figure 3E). On the other hand, following either chronic or acute HU treatment, the percentage of EdU-positive mycATRΔC-/- cells was significantly reduced compared with mycATR cells (Figure 3E). Interestingly, the nuclear signal of EdU positive cells did not show an overall difference between mycATR and mycATRΔC-/- cells, suggesting that once DNA replication restarts, the level of DNA synthesis is similar in both cell lines (Figure S3). Together, these data demonstrate that ATR is crucial for efficient DNA replication restart in *L. major*.

### ATR C-terminal deletion results in nuclear DNA damage

Our results above suggest that loss of ATR’s C-terminus coincides with cell cycle defects and enhanced genome instability. To test this interpretation, we next examined the levels of γH2A, a DNA damage marker in trypanosomatids. mycATR and mycATRΔC-/- cells were harvested in the absence of replication stress or following exposure to chronic and acute HU treatment as described above. After treatment, whole cell extracts were prepared at 0 and 5 h after chronic treatment, and 0 and 4 h after acute treatment and the levels of γH2A assessed by western blotting. γH2A signal was detected using anti-*T. brucei* γH2A antiserum that cross reacts with *Leishmania* γH2A (60). γH2A signal was quantified for all cell lines and intensity calculated relative to untreated mycATR cells (Figures 4A-D). After chronic HU treatment, a significant increase in γH2A levels was observed in mycATRΔC-/- cells when compared with mycATR cells (Figure 4C). However, after acute HU treatment, γH2A levels were similar in both cell lines (Figure 4D). These findings suggest that the loss of the C-terminus of ATR results results in enhanced genome damage when *L. major* cells are able to sustain DNA replication under chronic HU treatment. Furthermore, the G1/S synchronization observed after acute HU treatment caused increased DNA damage in mycATR cells, a level of damage that is not further aggravated in mycATRΔC-/- cells. Given an increase in the sub-G1 population after both chronic and acute HU treatment (Figure 3C), we characterised and quantified the types of aberrant cells presumed to arise based on the proportion of nuclear and kinetoplast staining in individual cells. Key events in the cell cycle of trypanosomatids include flagellum duplication, followed by the duplication and separation of the kinetoplast (mitochondrial DNA), and the duplication and separation of the nucleus before cell division (61,62). To assess these events, we stained the DNA of fixed mycATR and mycATRΔC-/- cells with DAPI after HU treatment and analysed the proportions of nuclei (N) and kinetoplasts (K) in individual cells, whose outline was defined using α-Tubulin (Figure 4E). While only 1-2% of mycATR cells had aberrant N-K configurations after both chronic and acute HU treatment (Figure 4F), approximately 20% of exhibited aberrant N-K configurations after HU treatments. The majority of these aberrant cells were classified as 0N1K, meaning they retained kinetoplast staining but lacked nuclear staining. These findings reveal that aberrant cell division, likely leading to the altered DNA content as seen by sub-G1 staining in FACS analysis (Figure 3C), arises after HU treatment in mycATRΔC-/- cells, and reiterates that ATR plays a role in *L. major* genome homeostasis after DNA replicative stress.

**Figure 4.**
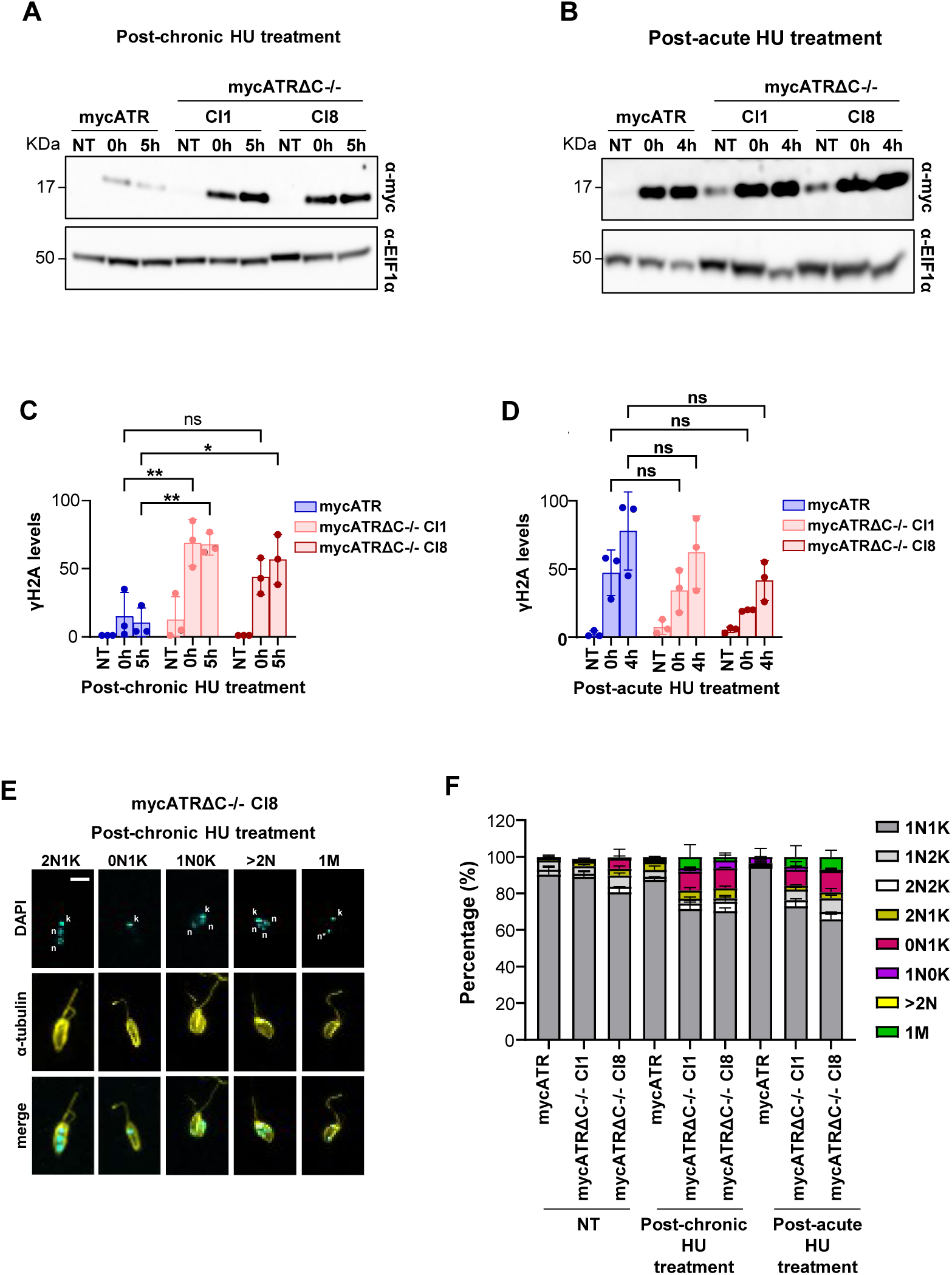
Deletion of the C-terminus of ATR leads to DNA damage and an increase of aberrant cells. (**A - B**) Western analysis of whole cell extract of each of the time points of Fig.3C after chronic and acute HU treatment; the blort was probed with α-γH2A antiserum; α-EIF1α was used as a loading control (predicted protein sizes are indicated, kDa). (**C – D**) Quantification of signal from the western blot showed in (A and B) using ImageJ software; statistical analysis were ised Graph Prism (Error bars ± SD; n = 3 experiments, * p<0.05; ** p<0.005; *** p<0.001; **** p<0.0001, 2way ANOVA). (**E**) Representative images acquired in LSM880 Zeiss confocal microscope from Z stack images of mycATRΔC-/- (cl8) after release from chronic (5 hours) treatment. Cells were fixed with 3% formaldehyde, genomic DNA stained with DAPI (cyan), and α-tubulin was used to stain the cytoskeleton (yellow). Bar scale = 9 um. (**F**) Percentage of DNA content populations based on the proportion of Nucleus/Kinetoplast (n/k) in mycATR and mycATRΔC-/- cells untreated (NT) or after release from chronic (5 hours) or acute (4 hours) HU treatment. The populations were coloured according to the proportion of n/k as normal (1n1k, 1n2n, 2n2k) or aberrant (0n1k, 1n0k, 2n1k, with more than 2 nucleus (>2N), and presence of Micronucleus, M). Error bars ± SD, n = 2 (>120 cells counted/experiment).

### ATR is needed to maintain the DNA replication program in *Leishmania major*

*L. major* exhibits a unique DNA replication program in which smaller chromosomes replicate before larger ones (63). This pattern may be connected to observations that only a single locus per chromosome is consistently detected as a primary site of DNA replication initiation (64) as detected by MFA-seq and DNAscent (64,65). Given that ATR is important to ensure the completion of DNA replication under both normal and replication stress conditions, we conducted MFA-seq analysis on both mycATR and mycATRΔC-/- cells, either untreated or treated with HU (Figure 5A) to assess if ATR plays a role during *Leishmania* DNA replication. Our work above revealed that exposing mycATRΔC-/- cells to chronic replication stress conditions permits DNA replication to progress, while mycATRΔC-/- cells displayed a pronounced impairment in DNA replication after acute HU treatment (Figure 3C). Hence, we used chronic replicative stress to test for replication timing defects.

**Figure 5.**
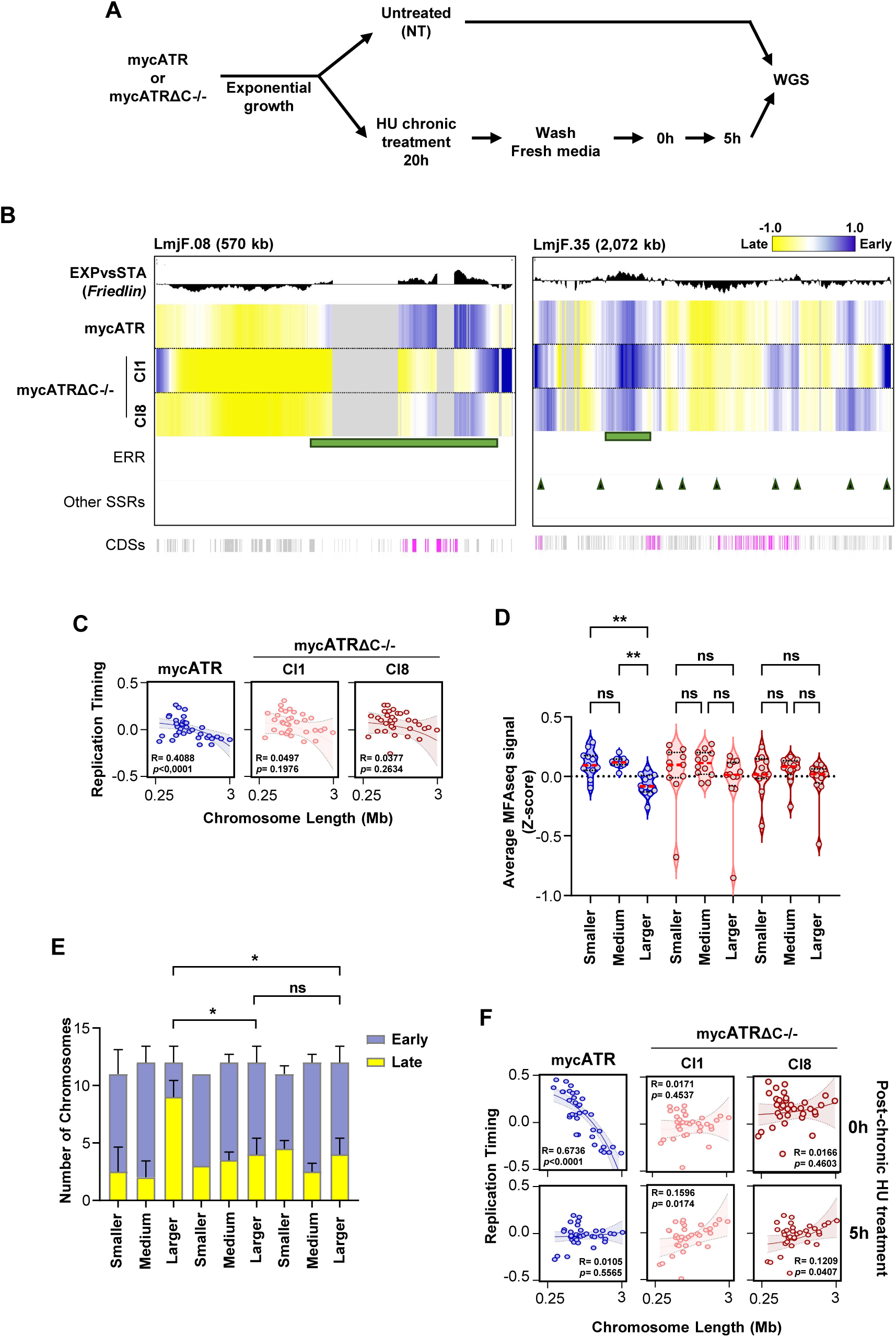
ATR is a determinant of the *L. major* DNA replication programme. (**A**) Schematic representation of the experimental design. mycATR or mycATRΔC-/- cells where left untreated (NT) or treat with 0.5mM of Hydroxyurea (HU) for 20 hours (Chronic). After treatment, cultures were washed and resuspend in fresh media. Time points were collected for sequence from NT cells 0, 5 hours after chronic treatment. Cells were passage 10 times after each condition and the DNA sequenced to evaluate long-term effects; DNA from stationary phase cells were sequenced to perform MFA-seq analysis (n = 2 experiments). (**B**) Representative snapshot of MFA-seq signal in chromosomes 08 and 35, showing DNA replication timing using exponentially untreated growing cells, mycATR and mycATRΔC-/- (cl1 and cl8), normalised with stationary cells; the signal represents the merge of the two replicates after normalisation; positive and negative values indicate early and late replicating regions, respectively; the top track represents a previous published MFA-seq from a wildtype *L. major* Friedlin (65); Early replication regions (ERR) are display in light green and the SSRs outside this regions displayed as arrow heads in dark green; the bottom track indicates annotated CDSs (gray: transcribed from left to right; pink: transcribed from right to left). (**C**) Linear regression analysis showing correlation between chromosome size and chromosome-averaged MFA-seq signal (DNA replication timing) of untreated mycATR and mycATRΔC-/- cells; R and P values are indicated at the right-top of each panel. (**D**) Quantification of the averaged MFA-seq in in untreated mycATR and mycATRΔC-/- cells. Chromosomes were sub-classified according with their length: Smaller (01 – 11 and 14), Medium (12, 13, 15 – 24) and Larger (25 – 36). (**E**) Quantification of the number of chromosomes that replicate according with MFA-seq average signal replicate Early (above 0) or Late (below 0) in untreated mycATR and mycATRΔC-/- (cl1 and cl8). Chromosomes were sub-classified as in 5D. (**F**) Linear regression analysis showing correlation between chromosome size and chromosome-averaged MFA-seq signal (DNA replication timing) after chronic HU treatment (0 and 5 hours) in mycATR and mycATRΔC-/-; R and P values are indicated at the right-top of each panel, rp1 and rp2 represent the two replicas.

To examine changes in DNA replication timing after ATR C-terminal deletion, we first plotted the MFA-seq profiles across individual chromosomes in the mycATRΔC-/- and mycATR cells (Figure 5B and S4), uncovering several key observations. First, the positive MFA-seq signal in mycATRΔC-/- cells appeared more broadly distributed across the larger chromosomes when compared with the MFA-seq signal from mycATR cells (where positive signal was predominantly concentrated at a single locus, as detected previously in wild type *L. major*; Figure 5B and S4). Second, this broader distribution of MFA-seq signal was not entirely consistent between the two mycATRΔC-/- clones, suggesting stochastic variation in the changed DNA replication program (Figure 5B and S4). Finally, there was a less clear change in MFA-seq distribution on the smaller chromosomes relative to the larger, with the exception of chromosome 8, which showed a reduced MFA-seq signal around late regions of the mycATRΔC-/- clones compared with mycATR (Figure 5B and S4).

Next, we assessed the correlation between DNA replication timing and chromosome size by linear regression analyses, using the average MFA-seq signal for each chromosome relative to its length (Figure 5C and S5). In the absence of HU treatment, mycATR cells exhibited a chromosome size-DNA replication timing profile similar to that seen wild type *L. major* (Figure 5C and S5) (63). Specifically, smaller and medium-sized chromosomes showed higher average MFA-seq signal (indicating earlier replication) compared with larger chromosomes, which exhibited lower average MFA-seq signal (indicating later replication). Remarkably, untreated mycATRΔC-/- cells showed no significant correlation between MFA-seq signal and chromosome size, suggesting a disrupted DNA replication program (Figure 5C and S5). This shift in replication timing is attributed to an increased MFA-seq signal (average > 0) (Figure 5D) in several larger chromosomes (n=7) in mycATRΔC-/-cells, compared with only a few (n=2) in mycATR cells (Figure 5E). Together, these findings indicates that loss of ATR’s C-terminus results in earlier replication of larger chromosomes (Figure 5D and 5E). Interestingly, chromosomes 8 and 31 exhibited significantly lower average MFA-seq signals compared with all other chromosomes, and this pattern remained unaffected by ATR mutation (Figure S5). This suggests that these chromosomes may have distinct replication dynamics. Taken together, our data indicate that deletion of ATR’s C-terminus disrupts the unique DNA replication timing of the *L. major* chromosomes, resulting in increased levels of DNA replication at several loci on the larger chromosomes. Interestingly, these changes in DNA replication programming do not appear to be associated with significant cell cycle defects (Figure 3C) or alterations in the proportion of replicating cells within the mutant population (Figure S1B).

To determine whether the changes in DNA replication timing observed in the ATR mutants were linked to DNA replication stress responses, we next analysed the replication timing profile of HU-treated mycATR and mycATRΔC-/- cells. We first performed linear regression analysis, again correlating the average MFA-seq signal of each chromosome with its length (Figure 5F). At 0 hours, when mycATR cells display an increased proportion of the population in early S-phase (Figure 3C), the chromosome-size-dependent replication timing profile observed in untreated cells was maintained: smaller chromosomes had higher average MFA-seq signals compared with larger chromosomes (Figure 5F). In contrast, after 5 hours, when most mycATR cells had traversed S-phase (Figure 3C), the MFA-seq signal was similar across both small and large chromosomes, indicating the completion of DNA replication (Figure 5F).

Loss of ATR’s C-terminus again disrupted the DNA replication timing profile (Figure 5F). As observed in untreated cells (Figure 5F), mycATRΔC-/- cells at 0 hours showed no correlation between average MFA-seq signal and chromosome length. However, 5 hours after chronic HU treatment, linear regression analysis revealed a reversal of the DNA replication program seen in unperturbed cells, with larger chromosomes showing higher average MFA-seq signals (Figure 5F). These findings suggest that the combination of ATR loss and chronic replication stress leads to a severe disruption of the normal DNA replication program typically observed in unperturbed *L. major* promastigote cells.

### ATR activity regulates DNA replication progression from early-S replicating loci and putative replication stress sites

To understand these alterations to DNA replication timing we describe above, we examined the MFA-seq signal across the genome under all conditions. The MFA-seq data revealed that as previously reported, DNA replication in chromosome predominantly initiates at a single locus, located at a strand switch region (SSR) where polycistronic transcription initiates and/or terminates, and proximal to the putative centromere (64–67). To investigate DNA replication around this ‘early replicating region’ (ERR), we generated metaplots of MFA-seq signal in mycATR and mycATRΔC-/- cells. We detected no quantifiable differences in MFA-seq signal between mycATR and mycATRΔC-/- cells at the 36 ERRs, whether analyzed collectively or separated by chromosome size (Figure S6A and S6B). This finding suggests that the altered DNA replication timing in the ATR mutants is not due to changes in replication initiation at these loci. We also generated metaplots of MFA-seq signal in mycATR and mycATRΔC-/- cells treated with HU. Unlike untreated cells, immediately after HU exposure, the metaplots showed a higher MFA-seq signal at the ERRs in mycATR cells compared to mycATRΔC-/- cells (Figure 6A and 6B). Furthermore, while the ERR signal decreased in mycATR cells after 5 hrs of HU treatment, it remained unaltered in mycATRΔC-/- cells (Figure 6A and 6B). These findings are consistent with the expected movement of DNA replication forks away from the ERRs from 0 to 5 hours (early S-phase to late S-phase; Figure 3C) in mycATR cells, and indicate that this progression is impaired in the absence of functional ATR in mycATRΔC-/- cells.

**Figure 6.**
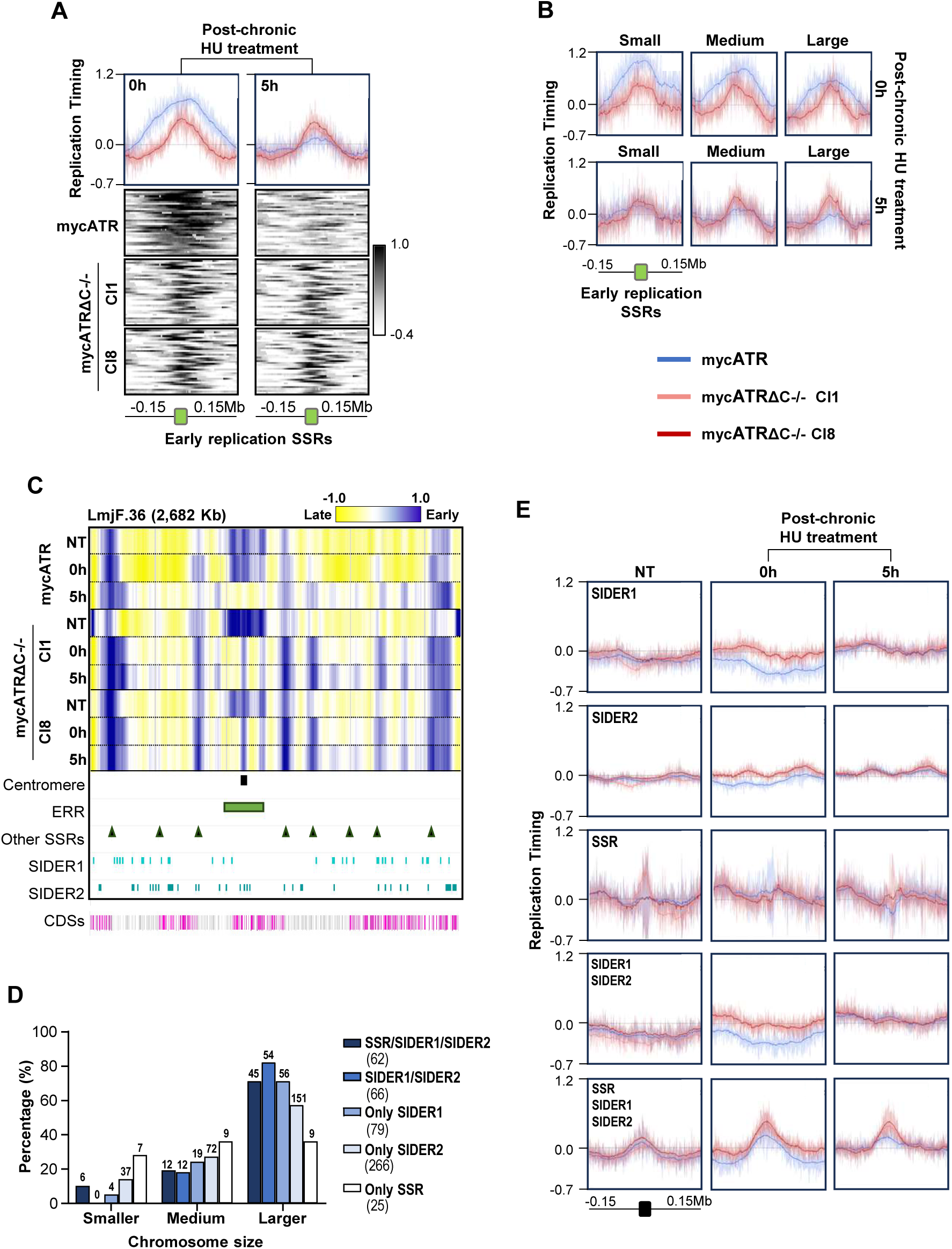
ATR activity is important for DNA replication progression both from the early replicating locus in each chromosome and around putative replicative stress regions. (**A**) Metaplots of global MFA-seq signal, in 0 and 5 hours after mycATR (blue) and mycATRΔC-/- (cl1 and cl8) cells be treated with chronic HU treatment relative to stationary cells ± 0.15 Mb from the centre of regions Early replication SSRs; the line indicate the mean and the light-coloured areas around the lines indicate the standard deviation resulted from the variation between the two experimental replicas (SD). Below: the colourmaps represent the MFA-seq signal in each early replication SSR in each sample. (**B**) Metaplots of global MFA-seq signal after subclassify the chromosomes according with each length: Smaller (01 – 11 and 14), Medium (12, 13, 15 – 24) and Larger (25 – 36). (**C**) Representative snapshot from MFA-seq signal in chromosome 36 showing the profile of DNA replication timing in mycATR and mycATRΔC-/- (cl1 and cl8) in untreated or after chronic HU treatment (0 and 5 hours). The displayed signal represents the merged signal from two replicas after the normalisation; positive and negative values indicate early and late replicating regions, respectively; KKT1 position is indicated in black, ERR in light green, SSRs outside ERR as arrow heads in dark green, SIDER1 in cyan and SIDER2 in dark cyan; the bottom track indicates annotated CDSs (gray: transcribed from left to right; pink: transcribed from right to left). (**D**) Graphic showing the percentage and number of SSR/SIDER1/SIDER2, SIDER1/SIDER2, or each of the three alone in each category of chromosome size (smaller, medium and larger). (**E**) Metaplots of global MFA-seq signal of mycATR and mycATRΔC-/- (cl1 and cl8) after chronic HU treatment (0 and 5 hours) around the regions described in (6D bottom) centred with ± 0.15 Mb from upstream and downstream of the start and end point of each region respectively.

Since the observed impairment of DNA replication progression from the ERRs in the absence of functional ATR and under low levels of HU cannot explain the reversal of the DNA replication timing programme, we next examined MFA-seq mapping across entire chromosomes (Figure 6C and S7). The impaired progression of DNA replication around the ERR in HU-treated mycATRΔC-/- cells compared with mycATR cells was readily apparent, as evidenced by a narrower positive MFAs-seq signal in the former. In addition, the profiles revealed the emergence of positive MFA-seq signal at several non-ERR sites in each chromosome after HU treatment. These new signals were more widespread after 5 hours of HU treatment than at 0 hours, and were more abundant in the mycATRΔC-/- cells than in mycATR cells (Figure 6C and S7). Visual inspection of the MFA-seq profiles revealed that, while some of these ‘new’ MFA-seq signals coincided with non-ERR SSRs, many did not. Thus, although predicted sites of RNA polymerase loading and unloading are known to be key locations of DNA replication stress in *L. major* (47), and given that collisions between the transcription and DNA replication machineries are a well-established trigger of ATR function (68), SSRs do not appear to be the sole locations of HU-induced MFA-seq signal accumulation. Two types of repetitive sequences that are part of short, interspersed degenerate retrotransposons (SIDERs), which are involved in posttranscriptional (SIDER1) and translational (SIDER2) gene expression regulation, cover almost 5% of the *L. major* genome(40,69–71). Similar to the SSRs, visual inspection suggested that some SIDER1 and SIDER2 sites in the genome might correlate with HU-induced MFA-seq signal (Figure 6C and S7), but not all. To test if there is an overlap between SSRs and SIDER elements with the HU-induced MFA-seq signals, we first examined the distribution of SIDER1 and SIDER2 across the genome (Figure 6D). SIDER1 elements (n=79) were less abundant than SIDER2 (n = 266) in non-ERRs. However, both types were more abundant on larger chromosomes. Notably, regions where either SIDER1 or SIDER2 was located within 5 Kb of an SSR were particularly common on the larger chromosomes (Figure 6D). Given this distribution, we generated metaplots of the MFA-seq signal from mycATR and mycATRΔC-/- cells, both untreated or after HU chronic treatment, around all possible combinations of non-ERR SSRs and SIDERs (Figure 6E). These plots revealed that there was no detectable accumulation of MFA-seq signal at any of the three sequence features alone under the tested conditions, consistent with re-analysis of previous MFA-seq data in wild type *L. major* cells (65) (Figure S6C). However, a modest enrichment of the MFA-seq signal was observed only at sites where SSRs overlapped with either SIDER1 or SIDER2 in untreated mycATRΔC-/- and mycATR cells, and was most pronounced at divergent (*DIV*) and Head-to-Tail (*HT*) SSRs (Figure 6E, S6D and S6E). These HU-induced signals displayed a pronounced increase at 0 and 5 hours post-HU treatment and in mycATRΔC-/- cells (Figure 6E, S6D and S6E). These findings suggest DNA replication at new loci after ATR loss and chronic HU exposure, particularly at loci on larger chromosomes, which may explain the disruption of chromosome size-dependent DNA replication timing.

### ATR is necessary for genome stability and variability in *Leishmania major*

Given that ATR partial deficiency exacerbates replication stress in *L. major* (47), we hypothesized that the loss of ATR function, along with the altered DNA replication program, might affect genome stability in *L. major*. To test this, we performed short-read Illumina whole genome sequencing of mycATRΔC-/- and mycATR cells exposed to chronic HU treatment and cultured for 10 passages (approximately 30 generations) (Figure 5A). We compared cells in the initial (P0) and 10^th^ passage (P10) to assess for changes in chromosome copy number (Copy Number Variations; CNV), a common indication of an untsable genome (Figure S8). mycATRΔC-/- cells showed a modest increase in CNV levels in several chromosomes in both conditions compared with mycATR cells (Figure S8A and S8B). Conversely, chromosomes 08 and 12 in untreated mycATRΔC-/- cells showed a decrease in the average CNV value after 10 passages when compared with mycATR cells (Figure S8A and S8B).

We also measured the formation of new single nucleotide polymorphisms (SNPs) and small insertions and deletions (InDels) after growth of mycATR and mycATRΔC-/- cells with and without HU treatment. This analysis revealed a pronounced chromosome size-dependent bias in accumulation of SNPs and InDels in mycATR cells, which became exacerbated by exposure to chronic replication stress (0.5 mM HU; Figure 7). Loss of ATR function in the mycATRΔC-/- cells appeared to have little effect on the accumulation of new SNPs in the absence of HU exposure, but SNP levels were notably reduced after HU exposure. InDels accumulation, in contrast, appeared to be reduced in all conditions by loss of ATR in mycATRΔC-/- cells (Figure 7). These data indicates that ATR acts to control CNV variation in *L. major* and its absence surprisingly suppresses localised mutations, suggesting there may be different paths that lead to genome variability in *L. major* and ATR function varies during these pathways.

**Figure 7.**
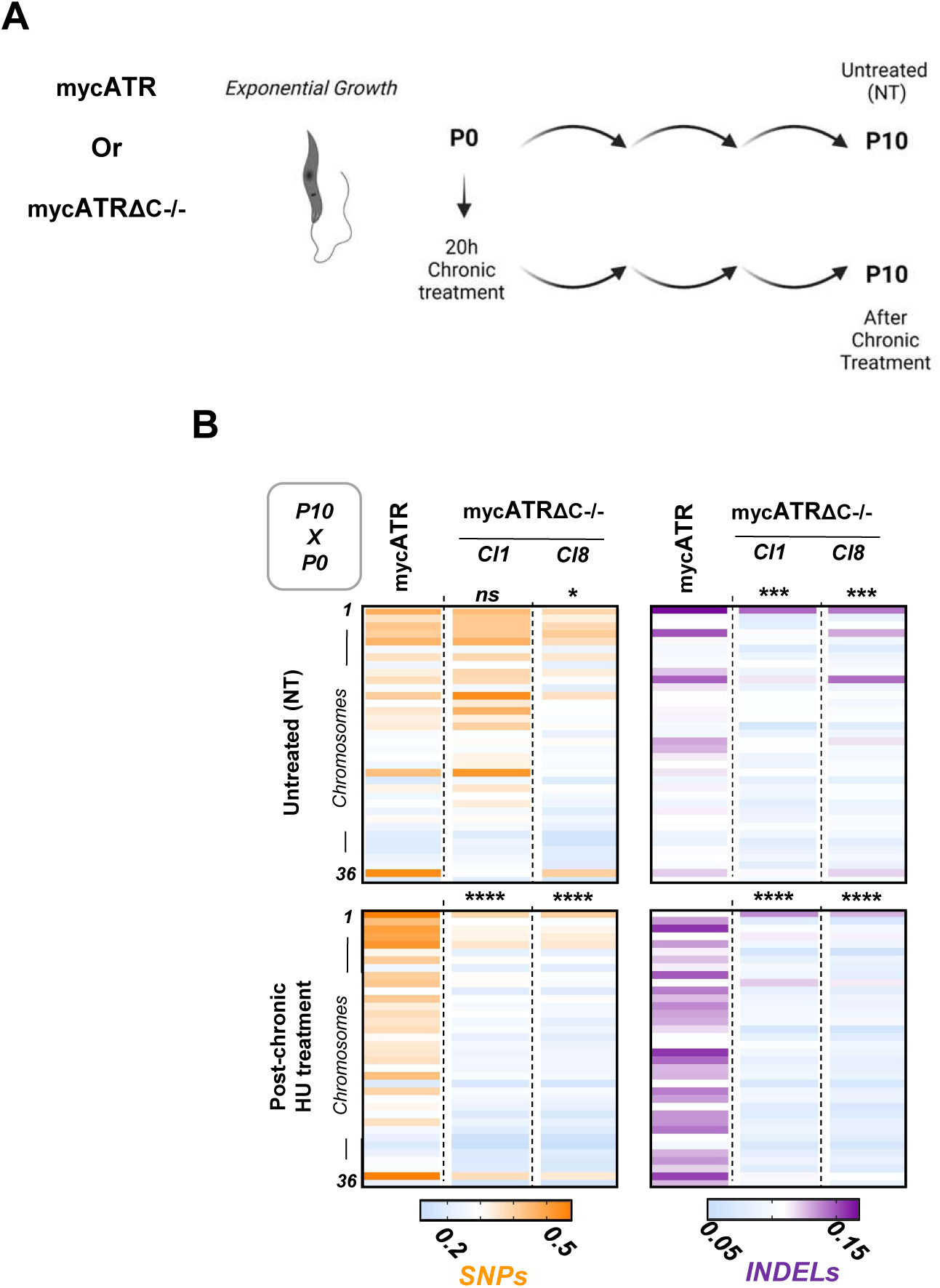
Replication program changes driven by ATR C-terminal deletion affect genome stability and variability in *Leishmania major*. Colourmap showing normalised density of new SNPs and InDels in each chromosome (ordered by size); new SNPs and InDels were calculated comparing p10/p0 for all untreated and HU treatment conditions and either cell lines mycATR and mycATRΔC-/- (cl1 and cl8).

## Discussion

In this study, we provide a genetic dissection of ATR function in *L. major* genome maintenance, building on previous studies that utilized ATR chemical inhibition (44,45). For reasons that remain unclear, we were unable to completely delete the *ATR* gene. Instead, we generated homozygous mutants lacking the C-terminus region of ATR, which includes the predicted kinase domain. The truncation led to a significant loss of ATR function, manifested at least in part through reduced protein abundance and loss of nuclear localisation. The evidence of such loss of function rests on several observations of the mycATRΔC-/- cells we generated: impaired growth *in vitro*, aberrant cell cycle progression, accumulation of ssDNA, and increased levels of nuclear damage seen as increased yH2A levels. We propose that these phenotypes reflect ATR’s conventional roles in managing DNA replication stress. This interpretation is consistent with the mutant’s heightened sensitivity to HU and the exacerbation of many of the above defects after HU treatment. Notably, our data also reveals a novel role for ATR in controlling the unusual DNA replication program of *L. major*, wherein smaller chromosomes are duplicated earlier than larger chromosomes (Figure 7).

ATR plays a central role in the cellular response to DNA damage, utilizing its kinase activity is used to regulate numerous repair pathways (2,3,25). One of the most well-studied pathways involves cell cycle control; for instance, ATR activates checkpoints in response to DNA damage, delaying progression through the S or G2 phases of the cell cycle (22,72). Our findings suggest that ATR plays a similar role in *L. major* and attribute this function to the C-terminus of the protein. Deletion of this region impairs the correct distribution of nuclear and kinetoplast (mitochondrial) DNA during cell division and disrupts cell cycle progression in the presence of HU. Similar effects are observed when ATR expression is knocked down in *T. brucei* (42,43); however, we cannot say here if these functions are solely dependent on kinase activity, as the truncated *L. major* ATR fails to efficiently localize to the nucleus. Nonetheless, this work contributes to the growing evidence that *L. major* ATR connects with wider factors needed to manage DNA replication progression. Previous studies showed that partial loss of Rad9 or Hus1 – components of the 9-1-1 complex – causes accumulation of cells in late S and G2 phases after acute HU treatment (73–76) and conditional deletion of Hus1 was shown to be lethal, resulting in the accumulation of ssDNA and γH2A – effects similar to those observed in mycATRΔC-/- cells (75). However, further investigation is needed to clarify *Leishmania*’s response to replication stress and ATR’s role in it. For instance, there are no reports of ATRIP or the ATR activators TOPBP1 or ETAA1 (77), and *L. major* 9-1-1 components appear to interact in distinct complexes during genome maintenance (78). Still, our data confirm ATR’s role in responding to replication stress across trypanosomatids.

An intriguing feature of *Leishmania* biology is its chromosome size-dependent DNA replication timing (63). This work reveals that ATR is a determinant of *L. major* DNA replication timing, since its mutation leads to earlier replication of the normally late-replicating large chromosomes. The basis of this change in timing is revealed by MFA-seq, which shows that unlike in wild type cells, where DNA replication initiation is predominantly detected in S-phase at just a single locus in each chromosome, MFA-seq peaks emerge at multiple sites across the chromosomes in the absence of functional ATR and, in particular, when the *L. major* ATR mutants are exposed to HU. One explanation for these new MFA-seq loci is that they represent normally suppressed origins, which are activated in the absence of ATR with and without excess replication stress caused by HU. Such a view would align with known functions of ATR in other eukaryotes, where licensed but not activated origins are controlled by ATR signalling (79), which acts as a negative regulator of origin firing and prevents inappropriate activation of origin firing in unperturbed cells (80), functions that are important to guarantee sufficient supply of DNA precursors and replication factors for optimal fork progression (80). So far, there has been little evidence for ‘dormant’ origins in trypanosomatids; in *T. brucei*, only one single putative site in one chromosome was found (81). However, although all SSRs appear to bind one subunit of ORC, only ∼25% are activated during unperturbed growth (82), and so SSRs may be a likely location for dormant origins that would be used in response to replication stress. Indeed, during DNA replication stress, ATR in other eukaryotes supresses origin firing globally, but allows dormant origin to be fired locally, preventing problematic replication across the genome (83). This might be consistent with our observations, where MFA-seq signal increases in mycATR cells around the single ERR in each chromosome immediately after HU chronic treatment, with more limited MFA-seq signal increase across the chromosomes than is seen in mycATRΔC-/- cells (Figure 6A and 6B). However, our data do not reveal the SSRs in *L. major* to be ubiquitous sites of increased MFA-seq signal after HU exposure, as might be predicted from ORC localisation in *T. brucei*. Instead, the major MFA-seq signals we see emerging after ATR mutation and HU exposure are where an SSR and a SIDER element are in proximity. While it remains possible that these loci are dormant origins, as these may untested sites of ORC recruitment, an alternative explanation is that these are genomic loci where DNA replication is especially prone to stalling, and this is exacerbated by the loss of ATR and exposure to HU. Hence, what we observe through MFA-seq may not be dormant origin activation, but sites of predominant DNA replication restart. This explanation would be consistent with two observations. First, the abundance of SIDERs in *Leishmania* has been associated with genome variation (30,70), consistent with them being problematic sequences for DNA replication. Indeed, it is possible that this is especially true around SSRs, as these are sites of RNA Polymerase loading and unloading, and hence SIDER proximity may increase clashes between transcription and DNA replication machineries. Second, the change in DNA replication timing we demonstrate here appears not to be limited to ATR function, since it is also seen after loss of RNase H1 or RAD51 (41,63), neither of which are known to directly coordinate origin activity: RNase H1 acts on RNA-DNA hybrids, which can form at sites of replication-transcription clashes, and RAD51 is the catalyst of DNA damage repair by homologous recombination, including at stalled replication forks (84–86). Thus, the normal course of DNA replication programming in *L. major* appears to be influenced by a range of activities that influence the effective completion of genome duplication.

Clarification of which of the above scenarios explain the change in DNA replication timing in *L. major* ATR mutants could come from examining the roles of a wider range of the parasite’s DNA repair factors in DNA replication timing. For instance, an increase of cells with aberrant DNA content is also observed after the conditional deletion of HUS1, indicating intersection of 9-1-1 and ATR activities (75). In fact, chromatin immunoprecipitation analysis of RPA1 and RAD9 showed that both ATR pathways factors accumulate around SSRs in unperturbed or during replicative stress cells (47). It may also be valuable to examine the effects of ATM loss since this DNA damage kinase acts on DNA double strand breaks and directs recruitment of RAD51.

In this study, we reveal that deletion of ATR’s C-terminus leads to lowered accumulation of mutagenic markers (SNP and InDels) after several rounds of cell multiplication, including in conditions of replication stress, suggesting a link between DNA replication and genome variation. Whether this is due to the change we observe in the normal pattern of *L. major* DNA replication timing is currently unclear, but it has been demonstrated that loci where DNA replication initiates are prone to accumulate SNPs (87,88), and, indeed, the *L. major* ATR mutant struggles to progress/re-start replication after replicative stress. Furthermore, CNV analysis showed that chromosomes 08 and 12 displayed the most substantial changes in content and were both notably distinct from the wider chromosomes in replication timing and showed low levels of MFA-seq signal increase after replicative stress. Though we do not know why these chromosomes differ, these data further suggest that deviation from the novel chromosome-size dependent pattern of DNA replication may have implications for stability. In conclusion, this work adds to the emerging picture of an intricate connection between DNA replication programming and genome variation in *Leishmania*.

## Methods

### Parasite culture and generation of ATR cell lines

Promastigotes derived from Leishmania major strain LT252 (MHOM/IR/1983/IR) were cultured at 26 °C in HOMEM medium supplemented with 10% heat-inactivated foetal bovine serum. Briefly, the primers were produced using the gene ID of ATR, LmjF.32.1460, on Leishgedit.net as described in(48). For each fragment amplified (sgRNA and donor fragment) five PCR reactions were performed as follows using Phusion polymerase (NEB, #M0530S) (98°C/1min, 98°C/30s, 60°C/30s, 72°C/45s or 1:45min, 72°C/5min, 35 cycles). All PCR reactions were precipitated and pooled as follows: Add ETOH 100% (cold) 250ul; 3M Sodium acetate 10ul; Glycoblue 3ul; and store at-20 °C (minimum 30min). After, the DNA was centrifuged as before for 20min, then washed in ETOH 70% (cold): 500ul by centrifugation as before for 20 min. The pellet was air dried then resuspended in 20ul of MilliQ water. For each transfection, 10 μl of the precipitated DNA was used. For recombination transfection or double tagging, *5x105 L. major* Cas9 cells were resuspend in 100 μl of 1× Tb-BSF buffer and pulse using the X-100 program in the Amaxa Nucleofactor™ IIb (Lonza)(89). After transfection, cells were resuspended in 10 mL of fresh media with 20% of FBS and recovered for at least 12 hours, then the appropriate selection drug was added, and the cells plated on a 96-well plate on the dilution of 1:6 – 1:72 – 1:864 for ∼ 5 weeks. First to generate *^myc^*ATR and *^3M^*ATR, expressing cells were selected using 10 μg. mL^-1^ puromycin (*Pur*) and 20 μg.mL^-1^ hygromycin (*Hyg*). Homozygotes *^myc^*ATR cell lines were used in a second round of transfection to generate *^myc^*ATR*^ΔC-/-^*cell line, and expressing cells were selected using 10 μg. mL^-1^ puromycin (*Pur*), 20 μg.mL^-1^ hygromycin (*Hyg*) and 1 μg.mL^-1^ neomycin (*Neo*). Integration into the expected locus was confirmed by PCR analysis. **Tables S1** and **Table S2** detailed the cell lines and list of primers used on this study, respectively.

### Antibodies

Primary antibodies used in this study were α-myc clone 4A (Millipore, #05-724), α-EIF1α (Millipore, #05-235), α-BrdU clone B44 (BD Bioscience, #347580), α-tubulin Beta clone KMX-1 (Millipore, #MAB3408), α-γH2A (1:1000) from rabbit serum was previously described(60). α-Mouse IgG Alexa Fluor 488-conjugated (ThermoFisher, #A11001), goat α-Rabbit IgG IRDye 800CW-conjugated (Li-COR Biosciences) and goat α-Mouse IgG IRDye 680CW-conjugated (Li-COR Biosciences). was used as secondary antibody.

### Western Blotting

Whole cell extracts were prepared by collecting cells by centrifugation, washing with 1x PBS + 1x protein inhibitor (ThermoFisher, #S8830), resuspending in extraction buffer: 1xNuPAGE™ LDS Sample Buffer (ThermoFisher), 10% of 1x protein inhibitor (Thermo Fisher) and 5% β-mercaptoethanol; heating the samples to 95°C for 7 minutes. Whole cell extracts were resolved on 4-12% gradient Bis-Tris Protein Gels (ThermoFisher) and transferred to Polyvinylidene difluoride (PVDF) membranes (GE Life Sciences) for 90min, 25V. Membranes were first blocked with 5% non-fat dry milk dissolved in 1x PBS supplemented with 0.05% Tween-20 (PBS-T) for 1 h at room temperature. Next, membranes were probed with primary antibody: α-myc (1:5000), α-EIF1α (1:25000), α-yH2A (1:1000); overnight at cold room 4°C, diluted in PBS-T supplemented with 2% non-fat dry milk. After washing with PBS-T, membranes were incubated with HRPconjugated secondary antibodies in the same conditions as the primary antibodies. Finally, membranes were washed with PBS-T. To detect and visualize bands, membranes were incubated with ECL Prime Western Blotting Detection Reagent (GE Life Sciences) and exposed to detection in a SynGenne Pxi touch gel Imaging System.

### Immunofluorescence and Detection of cells in S phase

2×10^6^ of exponential growing cells were collected by centrifugation, washed in 1.0 ml of 1xPBS and resuspend in 500ul of 3.7% paraformaldehyde for 15min at room temperature. After fixation, cells were washed in 1x PBS and 30ul of the cell was adhered into poly-L-lysine coated slides. Next, cells were resuspended in 1xPBS supplemented with 0.7% TritonX-100 for 20 min. Primary antibody were use diluted in 1x PBS + 3% BSA in following dilution: α-myc (1:500), α-tubulin (1:1000) and α-BrdU (1:500); and 30ul of antibody solution was used for 1 hour. After washing with 1x PBS + 3% BSA, Secondary antibodies were diluted as primaries in a dilution of 1:1000 for 45min. DAPI with fluoromount-G (ThermoFisher, #00-4959-52) were used to stain the DNA. Images were acquired using Zeiss Elyra Super resolution microscope or Zeiss LSM880 confocal microscope or Leica DMi8 microscope. Further image processing was performed with ImageJ software. For DNA synthesis detection with EdU; 2×10^6^ of exponential growing cells were left untreated or after chronic and acute hydroxyurea (HU) treatment incubated with 150uM of EdU (ThermoFisher) for a pulse of 30min. Slide preparation was made as described above, however, instead primary antibody, to detect EdU we used a Click-IT reaction (Invitrogen, #C10339).

### S phase detection and cell cycle profile using flow cytometry

Cells were incubated with 150 mM IdU (ThermoFisher) for 30 min (S phase detection); and then fixed at -20°C with a mixture (7:3) of ethanol and 1x PBS for at least 16 hr (from this step: cell cycle profile). Next, cells were rinsed with washing buffer (1x PBS supplemented with 1% BSA) and the DNA denatured for 30 min with 2N HCL, followed by neutralisation with phosphate buffer (0.2 M Na2HPO4, 0.2 M KH2PO4, pH 7.4). Detection of incorporated IdU was performed with α-BrdU antibody (diluted in washing buffer supplemented with 0.2% Tween-20) for 1 hour at room temperature. After washing, cells were incubated with α-mouse secondary antibody conjugated with Alexa Fluor 488 (diluted in washing buffer supplemented with 0.2% Tween-20) for 1 hour at room temperature and then washed. Finally, cells were stained with 1xPBS supplemented with 10 mg.mL^-1^ Propidium Iodide (PI) and 10 mg.mL^-1^ RNAse A and filtered through a 35 mm nylon mesh. FACSCelesta (BD Biosciences) was used for data acquisition and FlowJo software for data analysis. Negative control (omission of α-BrdU antibody during IdU detection step) was included in each sample and used to draw gates to discriminate positive and negative events and at least 30000 cells were used to obtain the results.

### Replication timing profiling using Marker Frequency Analysis coupled with Illumina sequencing (MFA-seq)

Genomic DNA was extracted from exponentially growing, treated with chronic or acute HU treatment and stationary cells using DNeasy Blood & Tissue Kit (QIAGEN) and were sequenced as 75 nucleotide paired-end reads. All libraries were prepared using using QIAseq FX DNA Library Kit (QIAGEN) and were sequenced as 75 nucleotide paired-end reads. Sequencing was performed at BGI (https://gtech.bgi.com/bgi/login) in a DNBSEQ-G400 sequencing platform. The Galaxy web platform (usegalaxy.org)(90) was used for most of the downstream data processing. For quality control and removal of adaptors, FastQC (http://www.bioinformatics.babraham.ac.uk/projects/fastqc/) and trimomatic(91) were used, respectively. Trimmed reads were mapped to the reference genome (Leishmania major Friedlin v39, available at Tritrypdb - http://tritrypdb.org/tritrypdb/) using BWA-mem(92). BamCompare (DeepTools) was used to determine reads abundance from exponentially growing cells relative to the reads from stationary culture. Ratios was first calculated in 500bp consecutive windows using the reads counts method for normalisation. Raw ratios were further transformed into Z scores relative to the whole genome ratios average as calculated in 100 kb sliding windows. **Table S3** list all the samples used in both replicas.

### CNV average

To calculate CNV ratios, the sequence depth was calculated in 2kb bin genome in each of the BAM files. The total number of reads was normalised by the depth and the CNV ratios calculated dividing the *normalised reads of sample P10*/ *normalised reads of sample P0 + 1×10^-6^* to prevent division by zero. The average of each of chromosomes were then obtain calculating the CNV values of each bin in each chromosome.

### SNP density

SNPs and InDels relative to the reference genome were detected in *p10* of each cell line in each condition and *p0* (untreated) cells using *FreeBayes(93)*. Only those SNPs and InDels in regions with read depth of at least 5, with at least 2 supporting reads, and a map quality of 30 were considered. To better capture the genomic variability in the time frame of the experiments, variants present simultaneously in *p10* and *p0* cells were excluded from the analysis using VCF-VFC intersect function from VCFtools package(94).

### Graphs plots generation and statistical analysis

MFAseq signal snapshots were obtained from Integrative Genomes Viewer (IGV). Heatmaps and metaplots were generated with deepTools plotHeatmap and plotProfile tools, respectively, on Galaxy. The intensity for western blot assay and fluorescence image were measured using ImageJ. Graphs were generated and statistical analyses were performed using Prism software. One-way an 2way analysis of variance (ANOVA) was used when comparing more than two groups. Comparisons between the two groups were made with the student’s t-test. Figure legends include P-values, sample size, and experiments were repeated at least 2 times using biologically independent.

## Data Availability

The data for this study have been deposited in the European Nucleotide Archive (ENA) at EMBL-EBI under accession number PRJEB83661.

## Acknowledgments

We thank the Molecular and Cell Biology of FMRP/USP staff and the University of Glasgow Centre for Parasitology for the support during the development of this project. This work was supported by FAPESP [16/16454-9, 19/20731-6], the Wellcome Trust [224501/Z/21/Z], and the BBSRC [BB/N016165/1, BB/R017166/1, BB/W001101/1]. Figures 5A, 7A and the abstract figure were generated using Biorender.

## Figure Legends

**Supplementary Figure 1. Characterisation of ATR kinase mutants in *Leishmania major***. (**A**) Growth curve of the indicated cells cultivated in HOMEM medium; cells were seeded at ∼2x105 cells.ml-1 at day 0; growth profile was evaluated after cells were kept in culture for 6 days and the cell density was assessed every 24 hours, CC1 and Cas9T7 cells were used as control (Error bars ± SD; n = 2 experiments). (**B**) Representative pseudocolor plots from a flow cytometry analysis to detect DNA synthesis in the indicated cell lines; mycATR and mycATRΔC-/- (cl1 and cl8). cells were seeded at ∼2x105 cells.ml-1 at day 0; at the indicated time points an aliquot of each cell line was incubated with IdU for 30 min and IdU detected under denaturing conditions; 30,000 cells were analysed per sample; 1N and 2N indicate single and double DNA content; dashed red lines indicate the threshold used to discriminate negative from IdU-positive events; inset numbers indicate total percentage of IdU-positive events on the whole population (n=2 experiment).

**Supplementary Figure 2. The ATR C-terminus is vital for accumulation of the kinase in the nucleus.** (**A**) Schematic representation of the region where we predict a possible Nuclear Location Signal (NLS). Predicted ATR protein sequence was accessed through Tritryp.pdb, and the search for possible NLS was made using two available software (http://nls-mapper.iab.keio.ac.jp/cgi-bin/NLS_Mapper_y.cgi) and (https://nucpred.bioinfo.se/cgi-bin/single.cgi). (**B**) Representative images of the sub-cellular localization of mNeongreen signal which is expressed N-terminally in live mycATR cells; scale bar = 13 μm. Images were captured on a SP6 microscope (Leica). DNA shown in cyan, were stained with PBS and Hoescht solution for 10 min/ 27°C followed by image acquisition on the microscope; the mNeongreen signal is in yellow. (**C**) Representative images acquired in LSM880 Zeiss confocal microscope from Z stack images of 3MATR. Cells were fixed with 3% formaldehyde and the stained with α-myc in yellow, and the genomic DNA stained with DAPI in gray (Bar scale = 11 um).

**Supplementary Figure 3. Replication stress leads to high levels of ssDNA and cell cycle defects which impairs replication restart in mycATRΔC-/- cells.** EdU intensity (A.U.) from positive EdU mycATR or mycATRΔC-/- cells in each condition. An area was drawn around each EdU positive cells nuclei based on the DAPI stain and the EdU signal intensity was measured using ImageJ software, (Error bars ± SD; n = 2 experiments, * p<0.05; ** p<0.005; *** p<0.001, One-way ANOVA).

**Supplementary Figure 4. ATR is essential to maintain the replication program in *Leishmania major*.** Representative snapshot of MFA-seq signal in chromosome 36, showing the profile of DNA replication timing in mycATR and mycATRΔC-/- (cl1 and cl8) in untreated or after chronic HU treatment (0 and 5 hours). The data represents the merged signal from two replicates after the normalisation; positive and negative values indicate early and late replicating regions, respectively; KKT1 position is indicated in black, ERR in light green, SSRs outside ERR as arrow heads in dark green, SIDER1 in cyan and SIDER2 in dark cyan; the bottom track indicates annotated CDSs (gray: transcribed from left to right; pink: transcribed from right to left).

**Supplementary Figure 5. ATR is essential to maintain the replication program in Leishmania major.** Linear regression analysis showing correlation between chromosome size and chromosome-averaged MFA-seq signal (DNA replication timing) of untreated or after chronic HU treatment (0 and 5 hours) of mycATR and mycATRΔC-/- cells; R and P values are indicated at the right-top of each panel

**Supplementary Figure 6. ATR activity is important for DNA replication progression both from the early replicating locus in each chromosome and around putative replicative stress regions.** (**A**) Metaplot of the global MFA-seq signal over the ERR regions (Centre) with ± 0.15 Mb from upstream and downstream of the start and end point of each region respectively. The MFA-seq signal from a previous report(65) of a wildtype cells (L. major Friedlin) was used as control. Colourmap below represent the distribution of the signal in each ERR per sample. (**B**) Metaplots of global MFA-seq signal after subclassify the chromosomes according with each length: Smaller (01 – 11 and 14), Medium (12, 13, 15 – 24) and Larger (25 – 36). (**C**) MFA-seq signal from a previous report(65) of a wildtype cells (L. major Friedlin) plotted around the regions described in (6D bottom) centred with ± 0.15 Mb from upstream and downstream of the start and end point of each region respectively. (**D**) Graphic showing the percentage and number of SSR/SIDER1/SIDER2, or SSRs alone (Only SSR) in each type of Switch Strand Region (convergent, divergent and head-to-tail). (**E**) Metaplots of global MFA-seq signal of mycATR and mycATRΔC-/- (cl1 and cl8) after chronic HU treatment (0 and 5 hours) around the each type of SSR (*COVergent, DIVergent, HT head-to-tail*) that are in proximity with either SIDER1 or SIDER2 centred with ± 0.15 Mb from upstream and downstream of the start and end point of each region respectively.

**Supplementary Figure 7. ATR activity is important for DNA replication progression both from the early replicating locus in each chromosome and around putative replicative stress regions.** (**A**) Representative snapshot from MFA-seq signal of all chromosomes showing the profile of DNA replication timing in mycATR and mycATRΔC-/- (cl1 and cl8) in untreated or after chronic HU treatment (0 and 5 hours). The displayed signal represents the merged signal from two replicas after the normalisation; positive and negative values indicate early and late replicating regions, respectively; KKT1 position is indicated in black, ERR in light green, SSRs outside ERR as arrow heads in dark green, SIDER1 in cyan and SIDER2 in dark cyan; the bottom track indicates annotated CDSs (gray: transcribed from left to right; pink: transcribed from right to left). (**D**) Table showing the total number and the percentage in each chromosome sub-class of: Top, Swich Strand Regions (SSRs), and repetitive regions (SIDER1 and 2) inside each Early replication regions; Bottom (in rest of the genome) which represents the proximity of SSR and repetitive regions, percentage of regions with proximity of the types of repetitive regions, and where those factors are located alone.

**Supplementary Figure 8. Loss of the ATR C-terminus leads to CNV changes.** Genome-wide relative copy number variation (CNV) analysis; chromosomes are ordered by size; CNV is the average per chromosome of 2kb bin of the [ratio(normalised reads from p10 cells /normalised reads from p0 cells)], for all untreated and HU treatment conditions and either cell lines mycATR and mycATRΔC-/- (cl1 and cl8).

## Notes

https://www.ebi.ac.uk/ena/browser/view/PRJEB83661

### Competing Interest Statement

The authors have declared no competing interest.

