## Supplementary figures for "ATR, a DNA damage kinase, modulates DNA replication timing in *Leishmania major*"

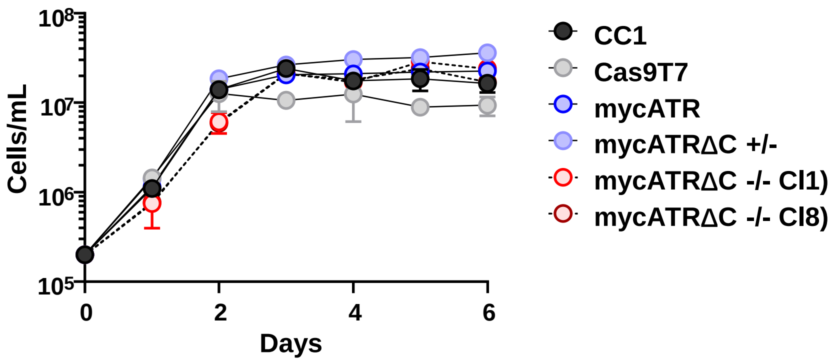
10^8
10^7
10^6
10^5
Cells/mL
0
2
4
6
Days
CC1
Cas9T7
mycATR
mycATRΔC +/-
mycATRΔC -/- Cl1)
mycATRΔC -/- Cl8)

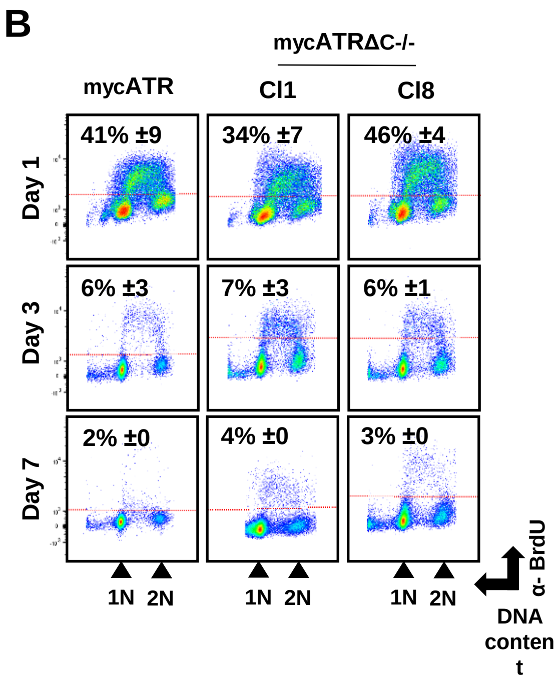
B
mycATRΔC-/-
mycATR
Cl1
Cl8
Day 1
41% ±9
34% ±7
46% ±4
Day 3
6% ±3
7% ±3
6% ±1
Day 7
2% ±0
4% ±0
3% ±0
1N 2N
1N 2N
1N 2N
α- BrdU
DNA conten t

**S1**

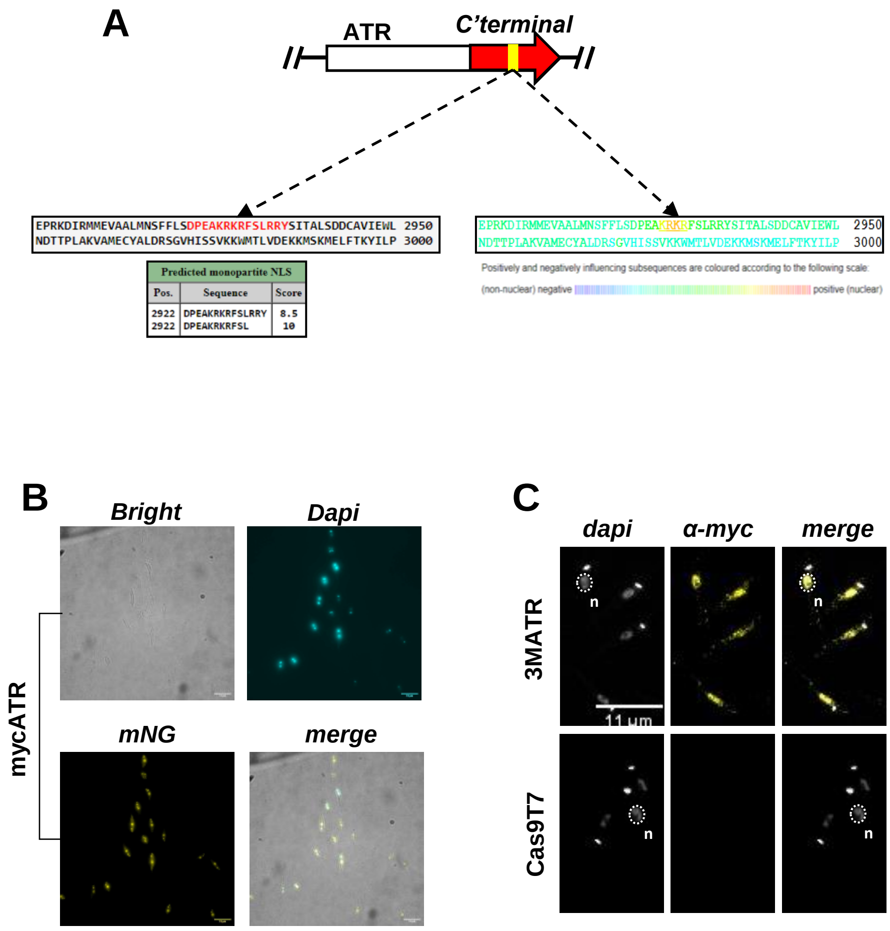
A
ATR
C'terminal
EPRKDIRMMEVAALMNSFFLSDPEAKRKRFSLRRYSITALSDDCAVIEWL 2950
NDTTPLAKVAMECYALDRSGVHISSVKKWMTLVDEKKMSKMELFTKYILP 3000
Predicted monopartite NLS
Pos. Sequence Score
2922 DPEAKRKRFSLRRY 8.5
2922 DPEAKRKRFSL 10
EPRKDIRMMEVAALMNSFFLSDPEAKRKRFSLRRYSITALSDDCAVIEWL 2950
NDTTPLAKVAMECYALDRSGVHISSVKKWMTLVDEKKMSKMELFTKYILP 3000
Positively and negatively influencing subsequences are coloured according to the following scale:
(non-nuclear) negative
positive (nuclear)
B
Bright
Dapi
mycATR
mNG
merge
C
dapi
α-myc
merge
3MATR
n
11 μm
Cas9T7

S2

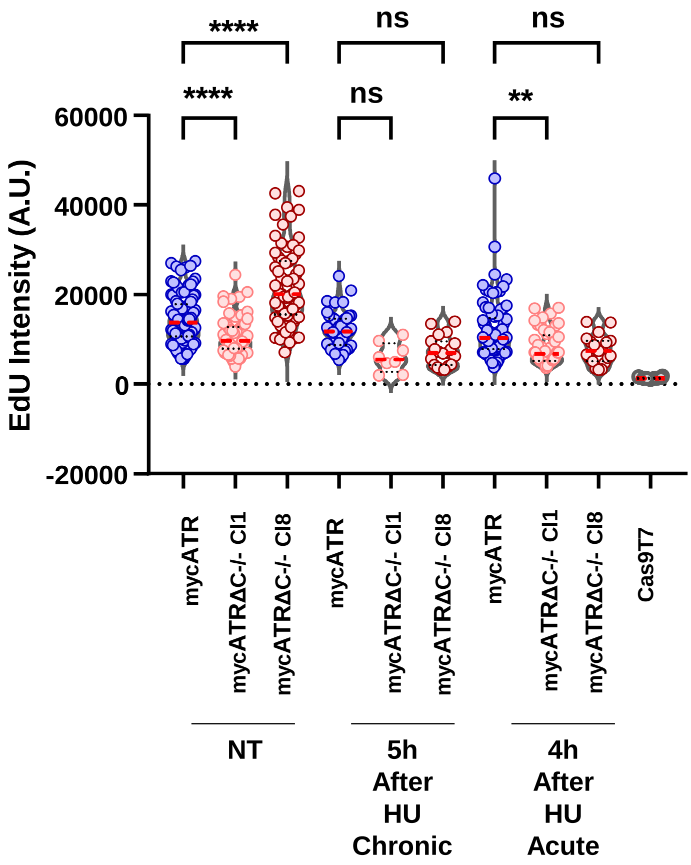
****
ns
ns
****
ns
**
EdU Intensity (A.U.)
60000
40000
20000
0
-20000
mycATR
mycATRΔC-/- Cl1
mycATRΔC-/- Cl8
mycATR
mycATRΔC-/- Cl1
mycATRΔC-/- Cl8
mycATR
mycATRΔC-/- Cl1
mycATRΔC-/- Cl8
Cas9T7
NT
5h After HU Chronic
4h After HU Acute

***S3***

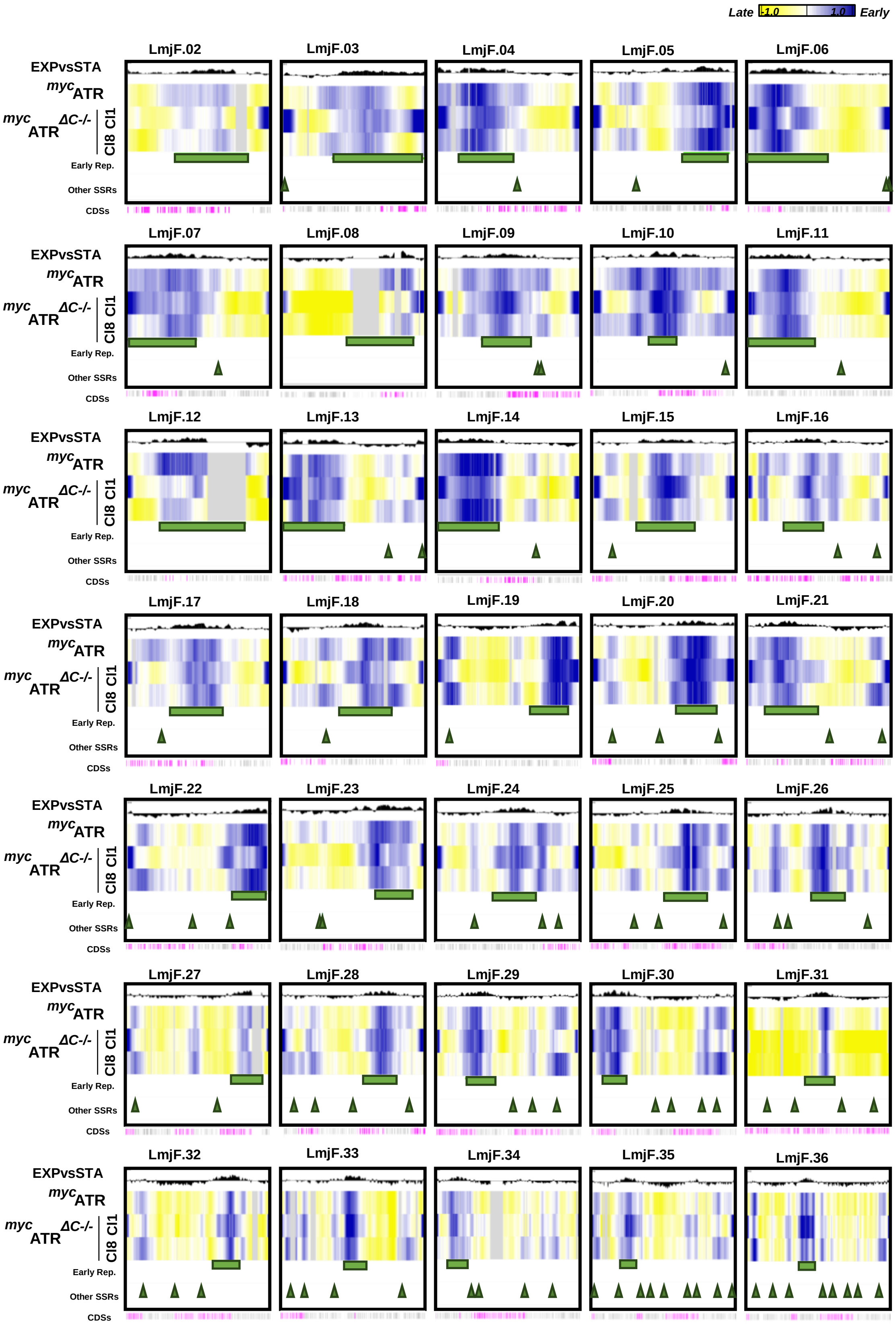
Late -1.0 1.0 Early
LmjF.02
LmjF.03
LmjF.04
LmjF.05
LmjF.06
LmjF.07
LmjF.08
LmjF.09
LmjF.10
LmjF.11
LmjF.12
LmjF.13
LmjF.14
LmjF.15
LmjF.16
LmjF.17
LmjF.18
LmjF.19
LmjF.20
LmjF.21
LmjF.22
LmjF.23
LmjF.24
LmjF.25
LmjF.26
LmjF.27
LmjF.28
LmjF.29
LmjF.30
LmjF.31
LmjF.32
LmjF.33
LmjF.34
LmjF.35
LmjF.36
EXPvsSTA
mycATR
mycATR ΔC-/-
Cl8 Cl1
Early Rep.
Other SSRs
CDSs

S4

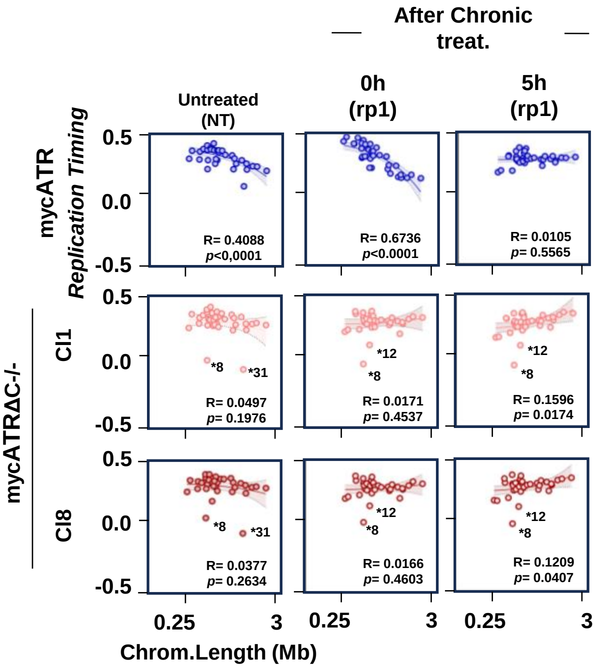
After Chronic treat.
Untreated (NT)
0h (rp1)
5h (rp1)
mycATR
Replication Timing
mycATRΔC-/-
Cl1
Cl8
0.5
0.0
-0.5
R= 0.4088 p<0,0001
R= 0.6736 p<0.0001
R= 0.0105 p= 0.5565
*8
*31
*12
R= 0.0497 p= 0.1976
R= 0.0171 p= 0.4537
R= 0.1596 p= 0.0174
R= 0.0377 p= 0.2634
R= 0.0166 p= 0.4603
R= 0.1209 p= 0.0407
0.25
3
Chrom.Length (Mb)

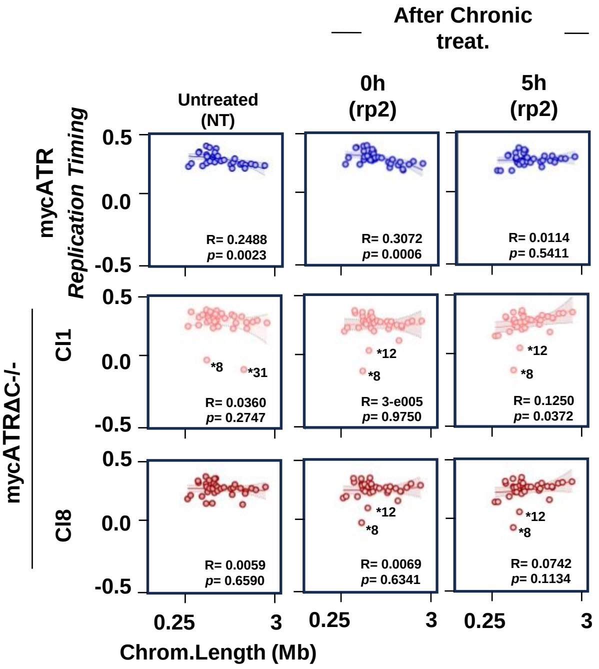
After Chronic treat.
Untreated (NT)
0h (rp2)
5h (rp2)
mycATR
Replication Timing
mycATRΔC-/-
Cl1
Cl8
0.5
0.0
-0.5
R= 0.2488 p= 0.0023
R= 0.3072 p= 0.0006
R= 0.0114 p= 0.5411
*8
*31
*12
R= 0.0360 p= 0.2747
R= 3-e005 p= 0.9750
R= 0.1250 p= 0.0372
R= 0.0059 p= 0.6590
R= 0.0069 p= 0.6341
R= 0.0742 p= 0.1134
0.25
3
Chrom.Length (Mb)

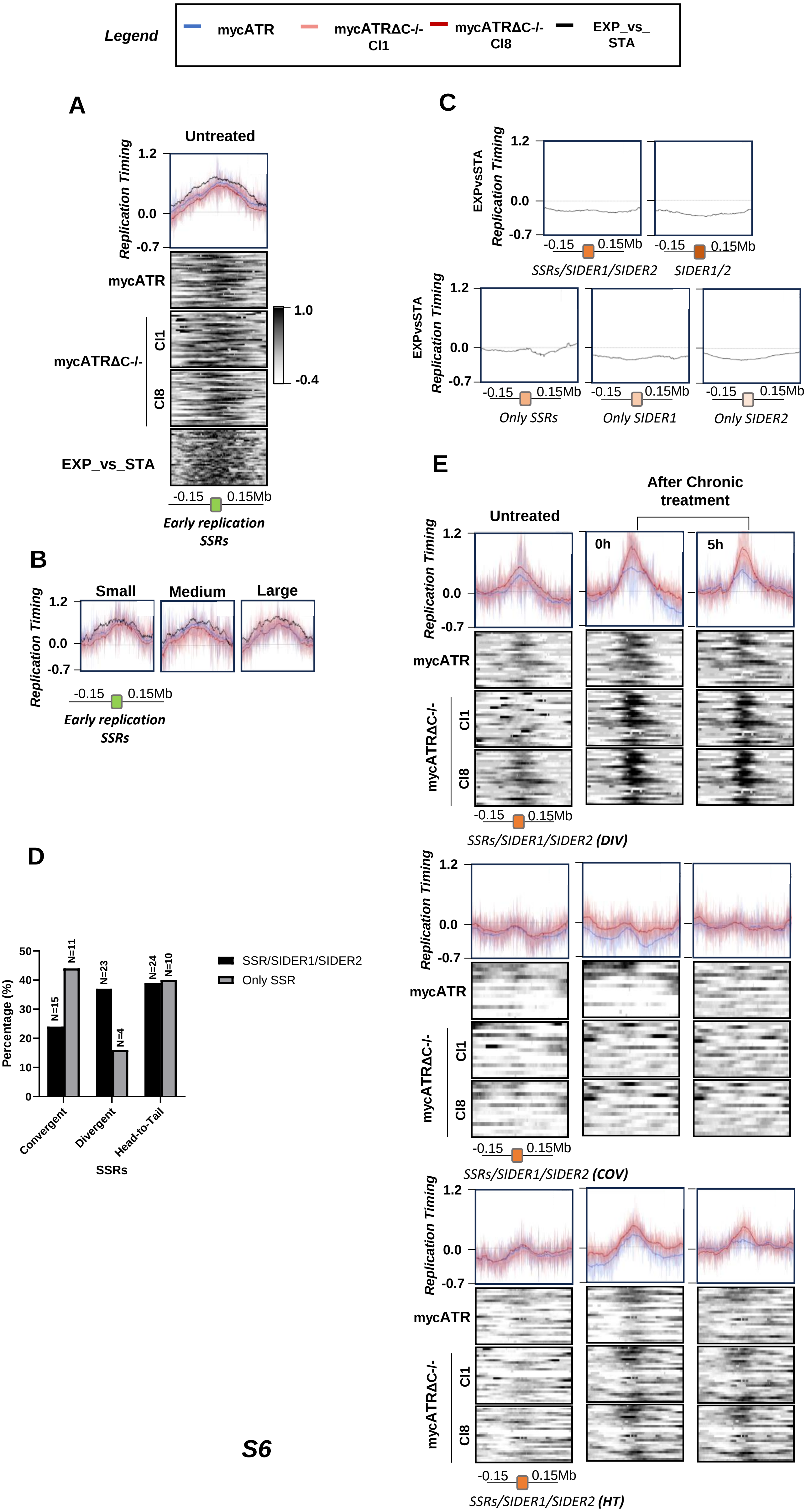
Legend
mycATR
mycATRΔC-/- Cl1
mycATRΔC-/- Cl8
EXP_vs_STA
A
Untreated
Replication Timing
1.2
0.0
-0.7
mycATR
mycATRΔC-/-
Cl1
Cl8
EXP_vs_STA
1.0
-0.4
-0.15
0.15Mb
Early replication SSRs
B
Small
Medium
Large
Replication Timing
1.2
0.0
-0.7
-0.15
0.15Mb
Early replication SSRs
C
EXPvsSTA
Replication Timing
1.2
0.0
-0.7
-0.15
0.15Mb
SSRs/SIDER1/SIDER2
SIDER1/2
Only SSRs
Only SIDER1
Only SIDER2
D
Percentage (%)
50
40
30
20
10
0
N=15
N=11
N=23
N=4
N=24
N=10
Convergent
Divergent
Head-to-Tail
SSRs
SSR/SIDER1/SIDER2
Only SSR
E
Untreated
After Chronic treatment
0h
5h
Replication Timing
1.2
0.0
-0.7
mycATR
mycATRΔC-/-
Cl1
Cl8
-0.15
0.15Mb
SSRs/SIDER1/SIDER2 (DIV)
SSRs/SIDER1/SIDER2 (COV)
SSRs/SIDER1/SIDER2 (HT)

S6

S7

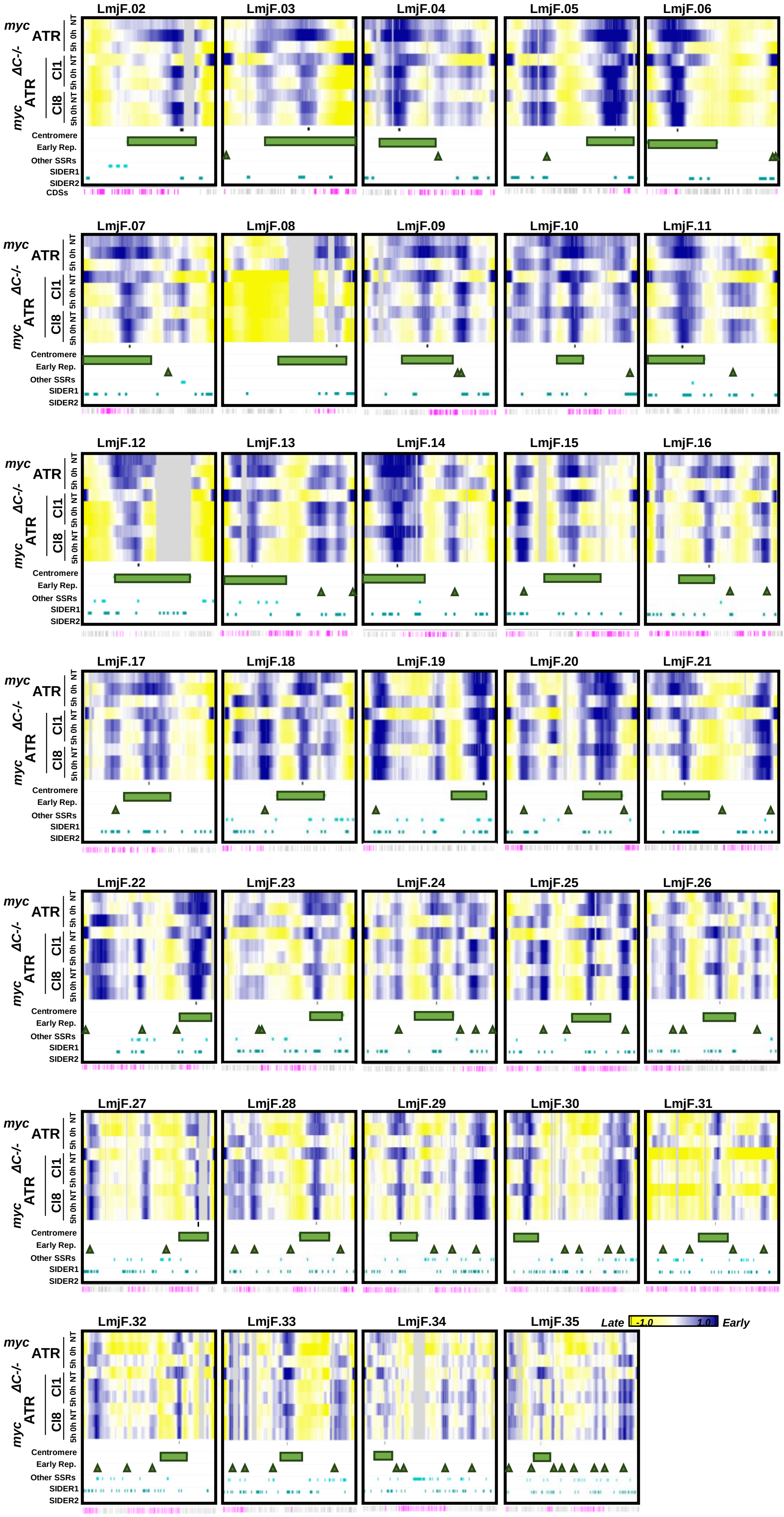

LmjF.02
LmjF.03
LmjF.04
LmjF.05
LmjF.06
LmjF.07
LmjF.08
LmjF.09
LmjF.10
LmjF.11
LmjF.12
LmjF.13
LmjF.14
LmjF.15
LmjF.16
LmjF.17
LmjF.18
LmjF.19
LmjF.20
LmjF.21
LmjF.22
LmjF.23
LmjF.24
LmjF.25
LmjF.26
LmjF.27
LmjF.28
LmjF.29
LmjF.30
LmjF.31
LmjF.32
LmjF.33
LmjF.34
LmjF.35
myc ATR
myc ΔC-/- ATR
Cl1
Cl8
NT
0h
5h
Centromere
Early Rep.
Other SSRs
SIDER1
SIDER2
CDSs
Late
-1.0
1.0
Early

**A**

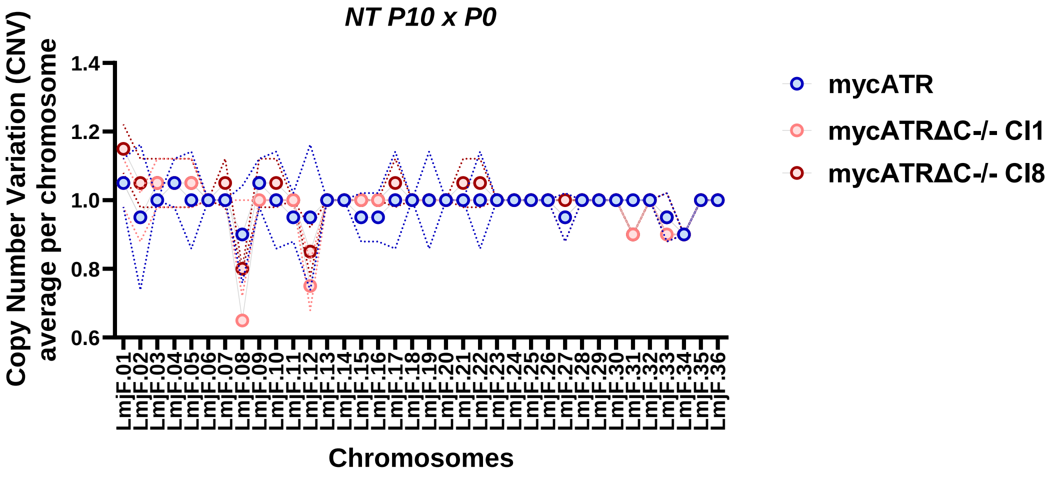

NT P10 x P0
Copy Number Variation (CNV)
average per chromosome
1.4
1.2
1.0
0.8
0.6
LmjF.01 LmjF.02 LmjF.03 LmjF.04 LmjF.05 LmjF.06 LmjF.07 LmjF.08 LmjF.09 LmjF.10 LmjF.11 LmjF.12 LmjF.13 LmjF.14 LmjF.15 LmjF.16 LmjF.17 LmjF.18 LmjF.19 LmjF.20 LmjF.21 LmjF.22 LmjF.23 LmjF.24 LmjF.25 LmjF.26 LmjF.27 LmjF.28 LmjF.29 LmjF.30 LmjF.31 LmjF.32 LmjF.33 LmjF.34 LmjF.35 LmjF.36
Chromosomes
mycATR
mycATRΔC-/- Cl1
mycATRΔC-/- Cl8


**B**

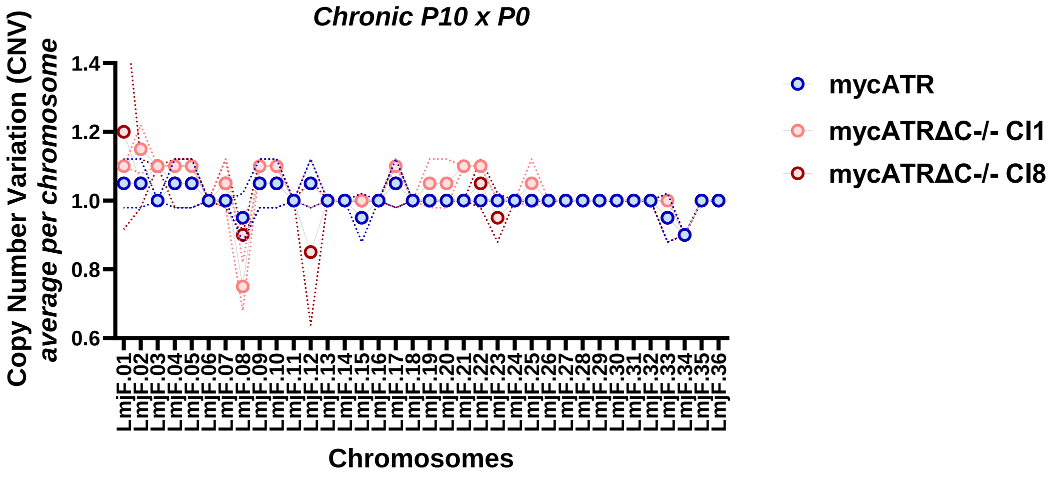

Chronic P10 x P0
Copy Number Variation (CNV)
average per chromosome
1.4
1.2
1.0
0.8
0.6
LmjF.01 LmjF.02 LmjF.03 LmjF.04 LmjF.05 LmjF.06 LmjF.07 LmjF.08 LmjF.09 LmjF.10 LmjF.11 LmjF.12 LmjF.13 LmjF.14 LmjF.15 LmjF.16 LmjF.17 LmjF.18 LmjF.19 LmjF.20 LmjF.21 LmjF.22 LmjF.23 LmjF.24 LmjF.25 LmjF.26 LmjF.27 LmjF.28 LmjF.29 LmjF.30 LmjF.31 LmjF.32 LmjF.33 LmjF.34 LmjF.35 LmjF.36
Chromosomes
mycATR
mycATRΔC-/- Cl1
mycATRΔC-/- Cl8

### Table 1 – cell lines on this study

| ***Name*** | ***Path*** | ***Select Marker*** | ***Description*** |
|---|---|---|---|
| **CC1** | **-** | **-** | Parental cell line |
| **Cas9T7** | ***>CC1*** | ***Hyg*** | Cell line expressing Cas9 endonuclease and T7 polimerase |
| ***myc*ATR** | ***>Cas9T7>CC1*** | ***Hyg, Pur*** | Cell line expressing Cas9 endonuclease and T7 polimerase and mNG+3myc tagging at ATR's N'terminal |
| ***3M*ATR** | ***>Cas9T7>CC1*** | ***Hyg, Pur*** | Cell line expressing Cas9 endonuclease and T7 polimerase and 3myc tagging at ATR's N'terminal |
| ***myc*ATR*ΔC+/-*** | ***>mycATR>Cas9T7>CC1*** | ***Hyg, Pur, Neo*** | Cell line expressing Cas9 endonuclease and T7 polimerase and mNG+3myc tagging at ATR's N'terminal and one allele deleted of ATR's C'terminal |
| ***myc*ATR*ΔC+/-*** | ***>mycATR>Cas9T7>CC1*** | ***Hyg, Pur, Neo*** | Cell line expressing Cas9 endonuclease and T7 polimerase and mNG+3myc tagging at ATR's N'terminal and both allele deleted of ATR's C'terminal |

***Table S1***

| Name | Sequence | Description |
| --- | --- | --- |
| a | TGTCACCTCTGTATTGGGCG | Fw upstream ATR gene |
| b | GCTGGTATTGCAGGAGGACA | Rv at ATR N'terminal gene |
| c | GCTGGACCGTTACATCTGGT | Fw at ATR gene |
| d | TACGGATGATGGCGCTACAC | Rv downstream ATR gene |
| e | GAAATTAATACGACTCACTATAGGATTGCTTCCCAGAGCAATGGGTTTTAGAGCTAGAAATAGC | 5' sgRNA ATR tagging |
| f | GAAATTAATACGACTCACTATAGGCAAGAGCAGACGGAGAGCCTGTTTTAGAGCTAGAAATAGC | 5'sgRNA ATR C'terminal deletion |
| g | GAAATTAATACGACTCACTATAGGGGACAAGCGCCTTGTCGTGCGTTTTAGAGCTAGAAATAGC | 3'sgRNA ATR C'terminal deletion |
| h | TTTTTGGCTCGATCGCGGAGCTCCTAGCCGGTATAATGCAGACCTGCTGC | Fw Donor mycATR and 3MATR tagging |
| i | GAGGCCCTCGTCGTCAGTGACAGCTTCCATACTACCCGATCCTGATCCAG | Rv Donor mycATR tagging |
| j | GAGGCCCTCGTCGTCAGTGACAGCTTCCATAGAACCGGAACCGGAACCAC | Rv Donor 3MATR tagging |
| k | CAGCTGCGGTGGACGCTGCTGCGCAATCGCGTATAATGCAGACCTGCTGC | Fw Donor mycATRΔC deletion |
| m | TAATACCCACAGGAGACACCAGTCCCGCACCCAATTTGAGAGACCTGTGC | Rv Donor mycATRΔC deletion |

***Table S2***

| ID | Sample | ID | Saample |
|---|---|---|---|
| GL4 | STA 1 | GL30 | mycatr_nt_p10_rp1 |
| GL5 | STA 2 | GL31 | cl8_acute_p10_rp1 |
| GL6 | STA 3 | GL32 | cl1_acute_p10_rp1 |
| GL7 | cl1_0h_acute_rp2 | GL33 | mycatr_chronic_5h_rp1 |
| GL8 | myatr_0h_acute_rp1 | GL34 | cl8_5h_chronic_rp1 |
| GL9 | cl1_0h_acute_rp1 | GL35 | cl8_5h_chronic_rp2 |
| GL10 | mycatr_0h_acute_rp2 | GL36 | cl1_5h_chronic_rp2 |
| GL11 | cl8_nt_rp1 | GL37 | cl1_chronic_5h_rp1 |
| GL12 | cl1_nt_rp1 | GL38 | mycatr_chronic_5h_rp2 |
| GL13 | cl1_nt_rp2 | GL39 | cl8_0h_chronic_rp2 |
| GL14 | mycatr_acute_p10_rp2 | GL40 | cl1_0h_chronic_rp1 |
| GL15 | mycatr_NT_rp2 | GL41 | cl8_rp1_0h_chronic |
| GL16 | cl1_nt_p10_rp1 | GL42 | mycatr_chronic_0h_rp1 |
| GL17 | mycatr_acute_p10_rp1 | GL43 | mycatr_0h_chronic_rp2 |
| GL18 | cl1_chronic_p10_rp2 | GL44 | cl1_chronic_0h_rp2 |
| GL19 | cl8_nt_rp2 | GL45 | cl1_acute_p10_rp2 |
| GL20 | cl8_p10_chronic_rp1 | GL46 | cl8_acute_p10_rp2 |
| GL21 | cl8_nt_p10_rp2 | GL47 | cl8_0h_rp2_acute |
| GL22 | mycatr_chronic_p10_rp1 | GL48 | cl8_acute_0h_rp1 |
| GL23 | mycatr_chronic_p10_rp2 | GL49 | cl1_acute_4h_rp1 |
| GL24 | mycatr_nt_rp1 | GL50 | cl8_acute_4h_rp1 |
| GL25 | cl1_nt_p10_rp2 | GL51 | mycatr_acute_4h_rp2 |
| GL26 | cl1_chronic_p10_rp1 | GL52 | cl1_acute_4h_rp2 |
| GL27 | mycatr_nt_p10_rp2 | GL53 | cl8_acute_4h_rp2 |
| GL28 | cl8_chronic_p10_rp2 | GL54 | mycatr_acute_4h_rp1 |
| GL29 | cl8_nt_p10_rp1 | | |

***Table S3***
